# A Multi-Omics Approach to Defining Target Organ Injury in Youth with Primary Hypertension

**DOI:** 10.1101/2024.06.17.599125

**Authors:** Kalyani Ananthamohan, Tammy M. Brady, Mohammed Arif, Stephen Daniels, Bonita Falkner, Michael Ferguson, Joseph T. Flynn, Coral Hanevold, Stephen R. Hooper, Julie Ingelfinger, Marc Lande, Lisa J. Martin, Kevin E. Meyers, Mark Mitsnefes, Bernard Rosner, Joshua A. Samuels, Gina Kuffel, Michael J. Zilliox, Richard C. Becker, Elaine M. Urbina, Sakthivel Sadayappan

## Abstract

**BACKGROUND:** Primary hypertension in childhood tracks into adulthood and may be associated with increased cardiovascular risk. Studies conducted in children and adolescents provide an opportunity to explore the early cardiovascular target organ injury (CV-TOI) in a population free from many of the co-morbid cardiovascular disease risk factors that confound studies in adults.

**METHODS:** Youths (n=132, mean age 15.8 years) were stratified by blood pressure (BP) as low, elevated, and high-BP and by left ventricular mass index (LVMI) as low- and high-LVMI. Systemic circulating RNA, miRNA, and methylation profiles in peripheral blood mononuclear cells and deep proteome profiles in serum were determined using high-throughput sequencing techniques.

**RESULTS:** *VASH1* gene expression was elevated in youths with high-BP with and without high-LVMI. *VASH1* expression levels positively correlated with systolic BP (r=0.3143, p=0.0034). The expression of hsa-miR-335-5p, one of the *VASH1-*predicted miRNAs, was downregulated in high-BP with high-LVMI youths and was inversely correlated with systolic BP (r=-0.1891, p=0.0489). *GSE1* hypermethylation, circulating PROZ upregulation (log_2_FC=0.61, p=0.0049 and log_2_FC=0.62, p=0.0064), and SOD3 downregulation (log_2_FC=-0.70, p=0.0042 and log_2_FC=-0.64, p=0.010) were observed in youths with elevated BP and high-BP with high-LVMI. Comparing the transcriptomic and proteomic profiles revealed elevated *HYAL1* levels in youths displaying high-BP and high-LVMI.

**CONCLUSIONS:** The findings are compatible with a novel blood pressure-associated mechanism that may occur through impaired angiogenesis and extracellular matrix degradation through dysregulation of Vasohibin-1 and Hyaluronidase1 was identified as a possible mediator of CV-TOI in youth with high-BP and suggests strategies for ameliorating TOI in adult-onset primary hypertension.

## INTRODUCTION

Primary Hypertension (PH) is a major risk factor for cardiovascular (CV) disease, affecting over one billion people globally.^1^ Chronic PH causes CV target organ injury (CV-TOI), including left ventricular hypertrophy (LVH), renal disease, and neurologic injury, which can result in heart failure, end-stage kidney disease, stroke, and cognitive dysfunction.^2–4^ Understanding the pathophysiology of PH and PH-associated CV-TOI is essential if novel diagnostic and therapeutic strategies are to be developed. Despite the worldwide prevalence, high mortality, and substantial healthcare expenditures^2^ related to PH, the actual mechanism(s) of PH and PH-associated CV-TOI is incompletely understood. Children and adolescents constitute an ideal population to study the origins of PH and its sequelae, as elevated blood pressure (BP) is detectable in the young,^5,6^ tracks into adulthood^7,8,9^ is increasing in prevalence^10^, and up to 40% of youth have evidence of LVH when hypertension is first diagnosed. ^11^ Moreover, while CV-TOI, such as LVH, is predictive of adverse hypertension-related outcomes in adults, the molecular associations of early PH in youth, including the evolution of hypertension-mediated CV injury in adolescence, remain unknown.^12^ Preliminary studies in animal models and patients suggest that antiangiogenic factors are central to pathological cardiac remodeling in cardiomyopathies^13–16^ and following ischemic injury.^17^ Tissue angiogenesis has been shown to have anti-fibrotic and cardioprotective effects among individuals with abnormal cardiac remodeling.^18–20^ Based on these studies, we explored correlations between the genetic and epigenetic regulation of circulating angiogenic factors and measures of CV-TOI in hypertensive youths.

The present study aimed to address the current knowledge gaps regarding PH-associated CV-TOI in youth by studying adolescent patients with and without elevated BP and LVH to identify clinical and molecular predictors of CV-TOI in circulation by integrating multi-omics data. Using transcriptomics data, we identified candidate genes and miRNAs involved in CV-TOI. Furthermore, we investigated circulating DNA methylation profiles and protein profiles to determine the potential pathways and their contributions to the CV-TOI process. Our results indicate that anti-angiogenic *VASH1* and its predicted miRNAs were dysregulated at the transcript level. Additionally, methylation and protein profiles identified candidate genes involved in inflammation, oxidative stress, and extracellular matrix degradation. Overall, we conclude that increased blood pressure is an early sign and contributes to the molecular changes involved in CV-TOI processes.

## RESULTS

### Circulating PBMC Blood Pressure Signatures in CV-TOI

To study the transcriptional dysregulation in circulation, we extracted RNA from the peripheral blood of study participants (high-BP+high-LVMI group, mid-BP, high-BP, low-BP+low-LVMI, and low-BP group) and then performed RNA-Seq. We identified numerous differentially regulated genes (DEGs) in the mid-BP (69 upregulated, 84 downregulated as compared to low-BP group) (Fig. 1A, Supplemental File S1), high-BP group (75 upregulated, 70 downregulated as compared to the low-BP group) (Fig. 1B, Supplemental File S1) and high-BP+high-LVMI group (4 upregulated, 6 downregulated as compared to the low BP+low LVMI) (Fig. 1C, Supplemental File S1) when considering exclusively a significance threshold of P-value ≤ 0.05 and log_2_fold change cutoff 0.5. Though this exploratory work identified DEGs, they failed to achieve any significance after the false discovery rate (FDR) correction. However, while comparing the DEGs with a significance threshold of P-value ≤ 0.05 without applying any fold change cutoff revealed common BP signatures. Three genes from the mid-BP group and 6 genes from the high-BP group were also upregulated in the high-BP+high-LVMI group (Fig. 1D). Similarly, 13 genes in the mid-BP group and 2 genes in the high-BP group were also downregulated in the high-BP+high-LVMI group (Fig. 1E). Using DisGenet, it is evident that these 24 genes identified by more than one comparison were known to be associated with hypertension or cardiovascular disease (Supplemental File S2). Thus, this initial analysis identified common BP signature genes dysregulated in high-BP study participants with and without high-LVMI.

**Fig. 1.**
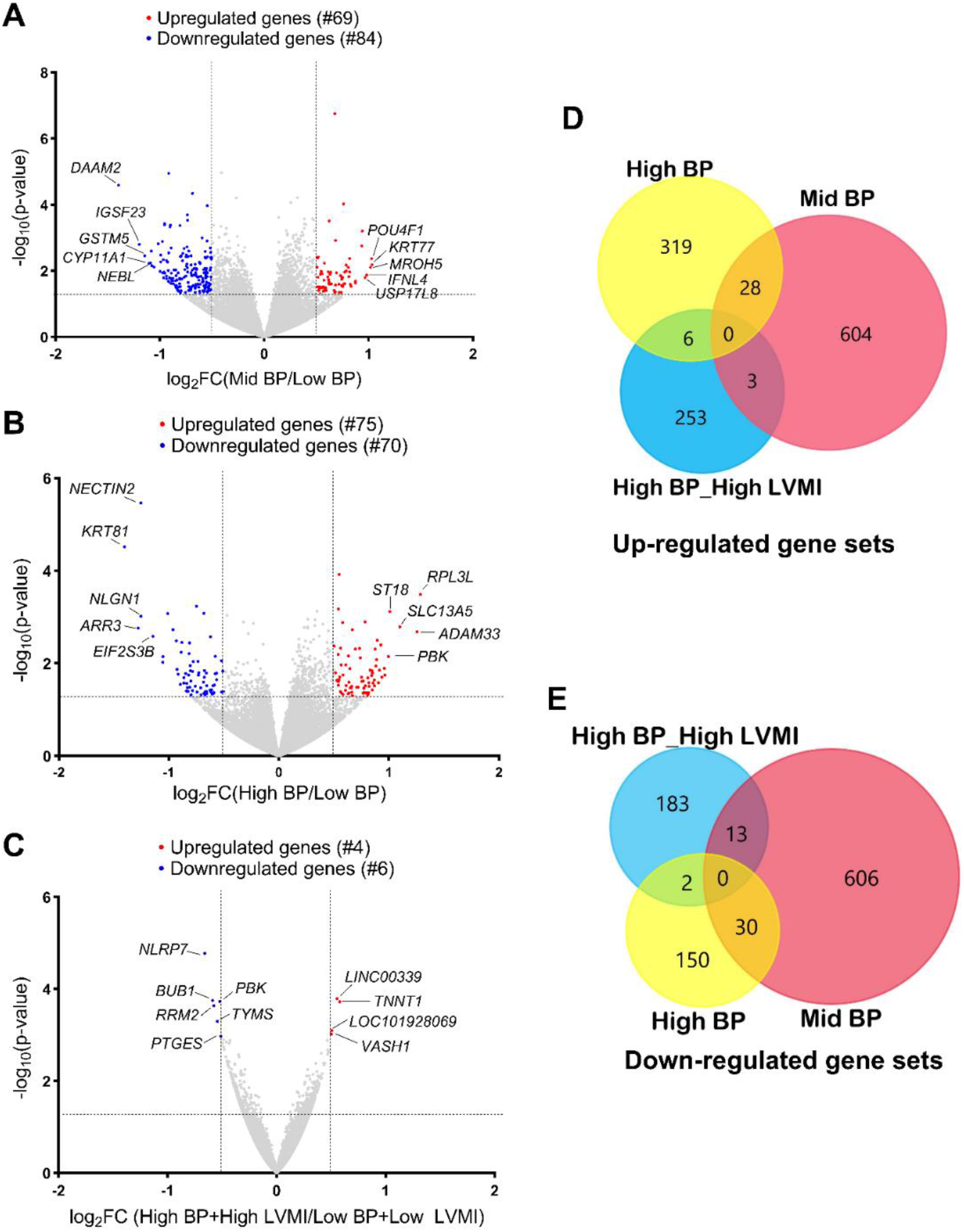
RNA expression pattern in PBMCs from SHIP AHOY participants with mid-BP, high-BP and high-BP+high-LVMI. Volcano Plot representing DEGs in mid-BP group **(A),** high-BP group **(B),** and high-BP+high-LVMI group **(C)** – each compared to their respective controls. RNAs with changes in expression and their p-value are determined by DEseq2 algorithm. The red color represents the increased gene expression, and the blue represents the diminished gene expression with p-value <0.05 and an absolute value of log_2_ fold change cut off 0.5. The top 5 up-regulated and down-regulated genes were labeled for each analysis. The upregulated and downregulated genes are compared among mid-BP, high-BP and high-BP+high-LVMI groups. **D-E,** Venn diagram showing the upregulated gene sets with a p-value <0.05 **(D)** and downregulated gene sets with a p-value <0.05 **(E)**. BP: Blood pressure; DEG: Differentially expressed genes; LVMI: Left Ventricular Mass Index; PBMCs: Peripheral blood mononuclear cells; SHIP AHOY: Study of High Blood Pressure in Pediatrics: Adult Hypertension Onset in Youth

### Transcriptomic Profiling Reveals Coordinated Regulation of Genes Linked to Blood Pressure

Our preliminary analysis identified the association between expression of 24 commonly dysregulated BP and cardiac candidate genes and CV-TOI in youth (Fig. 1D and 1E). The significant positive correlation between SBP PCT and LVMI (r=0.2804, p=0.0093, Supplemental Fig. S2), suggests that elevated blood pressure is associated with high-LVMI and may contribute to CV-TOI in hypertensive youth. Next, we analyzed the RNA expression values (FPKM) of the 24 DEGs with systolic BP percentile of participants in the low-BP, mid-BP, high-BP groups. This analysis found that *VASH1* (r=0.3143, p=0.0034, Fig. 2A), *EMP1* (r=0.222, p=0.0411, Fig. 2B), and *ST18* (r=0.2811, p=0.0092, Fig. 2C) were positively correlated with increasing SBP percentile, whereas *CACNB4* (r=-0.2783, p=0.009, Fig. 2D), *CTNNA2* (r=-0.2649, p=0.0143, Fig. 2E), and *CYP11A1* (r=-0.2379, p=0.0283, Fig. 2F), were negatively correlated with SBP percentile (Table 3). The correlation of these genes with LVMI displayed a similar trend only for *CTNNA2* and *CYP11A1* genes with significant negative correlation (Table. 3).

**Fig. 2.**
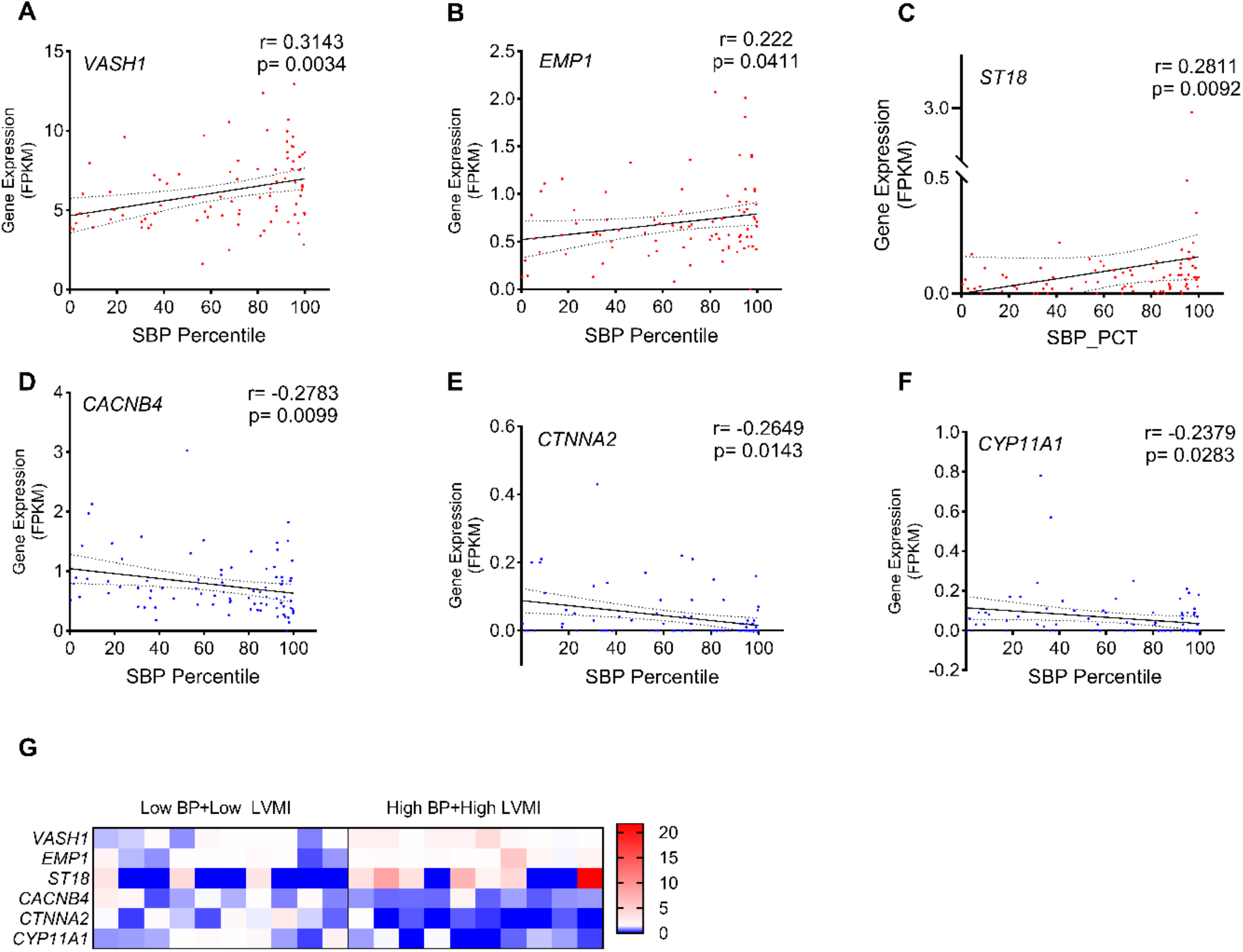
Distribution and correlation of LVMI and DEGs with systolic blood pressure percentile. Relation between SBP percentile and expression of *VASH1* **(A),** *EMP1* **(B)**, *ST18* **(C)**, *CACNB4* **(D)**, *CTNNA2* **(E)**, *CYP11A1* **(F)** among youth in the low-BP, mid-BP, and high-BP groups (n=85). These DEGs were selected for this correlation analysis from 24 candidates identified as common among the mid-BP group, high-BP group and high-BP+high-LVMI groups. Spearman correlation coefficients and their respective p-values were calculated. **(G)**, Expression of *VASH1*, *EMP1*, ST18, *CACNB4*, *CTNNA2*, and *CYP11A1* in low-BP+low LVMI group (n=10) and high-BP+high-LVMI group (n=10) are shown as a heat map for individual samples. The normalized counts for each gene are generated through an RNA Seq data analysis pipeline and all the values are normalized to the mean of their respective control group. DEG: Differentially expressed genes; FPKM: fragments per kilobase of exon per million mapped fragment values; LVMI: Left Ventricular Mass Index; SBP: systolic blood pressure.

**Table 1:**
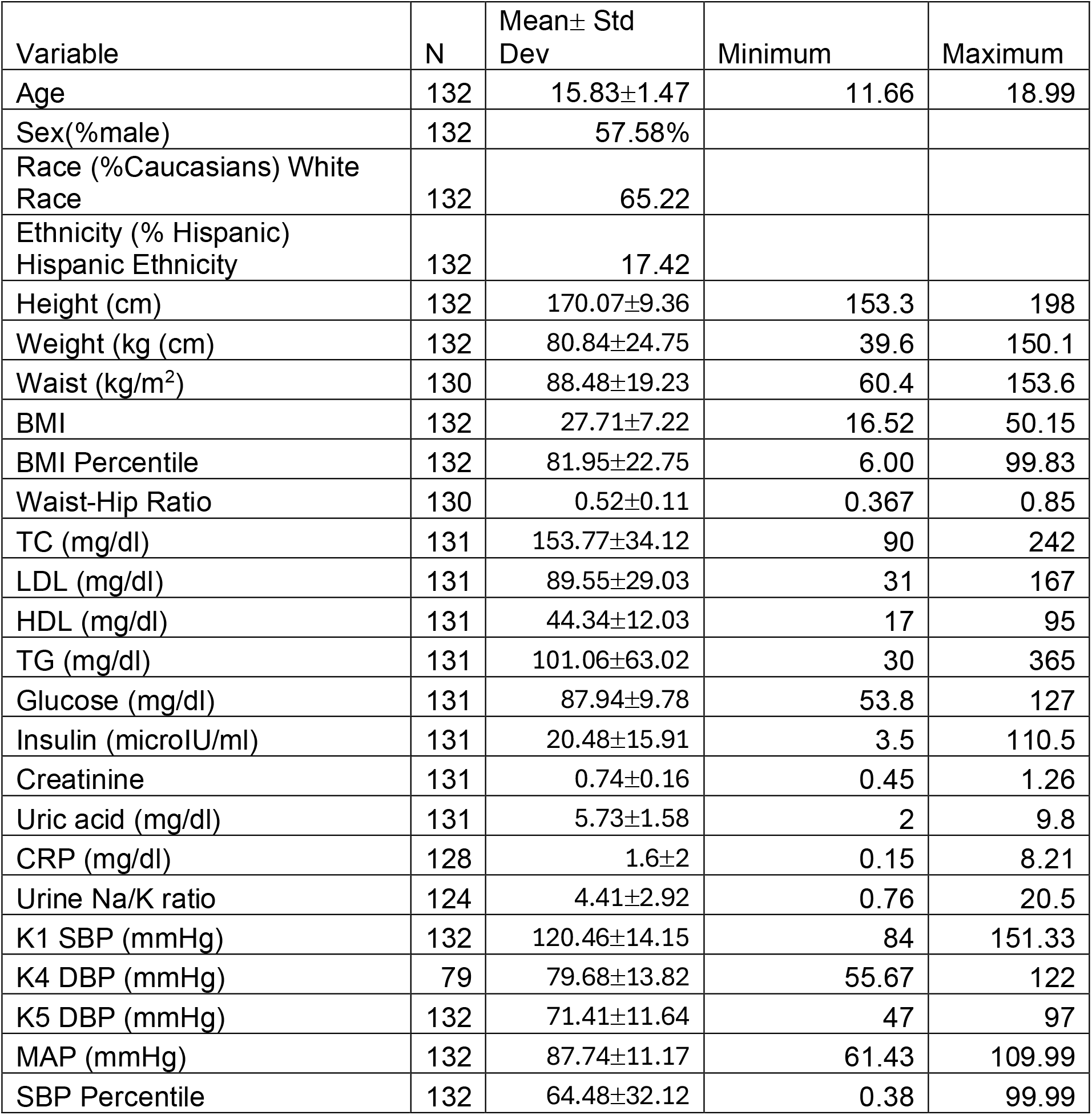
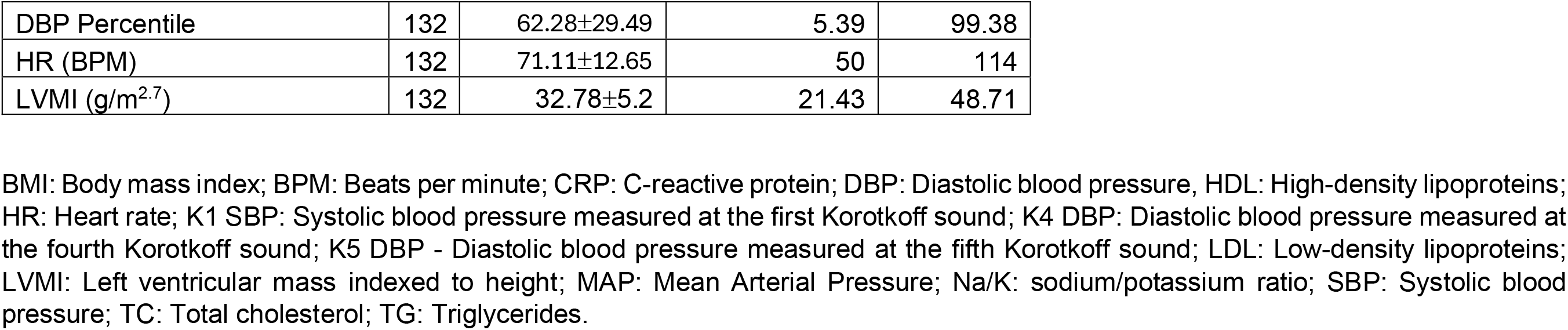
Description of the 132 Adolescents selected from SHIP AHOY study participants for the multi-omics study.

| Variable | N | Mean± Std Dev | Minimum | Maximum |
| --- | --- | --- | --- | --- |
| Age | 132 | 15.83±1.47 | 11.66 | 18.99 |
| Sex(%male) | 132 | 57.58% |  |  |
| Race (%Caucasians) White Race | 132 | 65.22 |  |  |
| Ethnicity (% Hispanic) Hispanic Ethnicity | 132 | 17.42 |  |  |
| Height (cm) | 132 | 170.07±9.36 | 153.3 | 198 |
| Weight (kg (cm) | 132 | 80.84±24.75 | 39.6 | 150.1 |
| Waist (kg/m<sup>2</sup>) | 130 | 88.48±19.23 | 60.4 | 153.6 |
| BMI | 132 | 27.71±7.22 | 16.52 | 50.15 |
| BMI Percentile | 132 | 81.95±22.75 | 6.00 | 99.83 |
| Waist-Hip Ratio | 130 | 0.52±0.11 | 0.367 | 0.85 |
| TC (mg/dl) | 131 | 153.77±34.12 | 90 | 242 |
| LDL (mg/dl) | 131 | 89.55±29.03 | 31 | 167 |
| HDL (mg/dl) | 131 | 44.34±12.03 | 17 | 95 |
| TG (mg/dl) | 131 | 101.06±63.02 | 30 | 365 |
| Glucose (mg/dl) | 131 | 87.94±9.78 | 53.8 | 127 |
| Insulin (microIU/ml) | 131 | 20.48±15.91 | 3.5 | 110.5 |
| Creatinine | 131 | 0.74±0.16 | 0.45 | 1.26 |
| Uric acid (mg/dl) | 131 | 5.73±1.58 | 2 | 9.8 |
| CRP (mg/dl) | 128 | 1.6±2 | 0.15 | 8.21 |
| Urine Na/K ratio | 124 | 4.41±2.92 | 0.76 | 20.5 |
| K1 SBP (mmHg) | 132 | 120.46±14.15 | 84 | 151.33 |
| K4 DBP (mmHg) | 79 | 79.68±13.82 | 55.67 | 122 |
| K5 DBP (mmHg) | 132 | 71.41±11.64 | 47 | 97 |
| MAP (mmHg) | 132 | 87.74±11.17 | 61.43 | 109.99 |
| SBP Percentile | 132 | 64.48±32.12 | 0.38 | 99.99 |
| DBP Percentile | 132 | 62.28±29.49 | 5.39 | 99.38 |
| HR (BPM) | 132 | 71.11±12.65 | 50 | 114 |
| LVMI (g/m<sup>2.7</sup>) | 132 | 32.78±5.2 | 21.43 | 48.71 |

BMI: Body mass index; BPM: Beats per minute; CRP: C-reactive protein; DBP: Diastolic blood pressure, HDL: High-density lipoproteins; HR: Heart rate; K1 SBP: Systolic blood pressure measured at the first Korotkoff sound; K4 DBP: Diastolic blood pressure measured at the fourth Korotkoff sound; K5 DBP - Diastolic blood pressure measured at the fifth Korotkoff sound; LDL: Low-density lipoproteins; LVMI: Left ventricular mass indexed to height; MAP: Mean Arterial Pressure; Na/K: sodium/potassium ratio; SBP: Systolic blood pressure; TC: Total cholesterol; TG: Triglycerides.

**Table 2:** Systolic blood pressure percentile and LVMI of SHIP AHOY participant stratified in different group.

| Variable<br>(Mean± S.D) | Low-BP + low-LVMI<br>N=10 | Low-BP<br>N=55 | Mid-BP<br>N=22 | High-BP<br>N=35 | High-BP + high-LVMI<br>N=10 |
| --- | --- | --- | --- | --- | --- |
| Systolic Blood<br>Pressure<br>Percentile | 27.5±21.05 | 41.28±23.25 | 85.1±3.09 | 95.5±3.02 | 94±4.75 |
| LVMI (g/m<sup>2.7</sup>) | 25.41±1.75 | 31.11±2.86 | 33.58±5.31 | 33.53±5.64 | 41.73±1.75 |

LVMI: Left ventricular mass indexed to height; Low BP indicates SBP > 75<sup>th</sup>% to <80<sup>th</sup>% , mid-BP indicates SBP 80<sup>th</sup>% - <90<sup>th</sup>% , high-BP indicates SBP ≥90<sup>th</sup>%

**Table 3.** Correlation of gene expression as determined from RNA seq data of PBMCs with cardiovascular disease risk factors.

| Genes (FPKM) | Systolic BP Percentile (n=85) |  | Left Ventricular Mass Index (g/m<sup>2.7</sup>) (n=85) |  |
| --- | --- | --- | --- | --- |
|  | Spearman r | p-value | Spearman r | p-value |
| *VASH1* | 0.3143 | **0.0034** | 0.1208 | 0.2708 |
| *EMP1* | 0.222 | **0.0411** | 0.07018 | 0.5233 |
| *ST18* | 0.2811 | **0.0092** | -0.03171 | 0.7733 |
| *CACNB4* | -0.2783 | **0.0099** | -0.0639 | 0.5612 |
| *CTNNA2* | -0.2649 | **0.0143** | **-0.2152** | **0.048** |
| *CYP11A1* | -0.2379 | **0.0283** | **-0.2691** | **0.0128** |

FPKM: fragments per kilobase of exon per million mapped fragment values; PBMC: peripheral blood mononuclear cells

This correlation analysis suggests that circulating PBMCs have distinct molecular pathways governing BP and LVMI in these subjects, which includes anti-angiogenesis (*VASH1*), C2H2C-type zinc finger transcription factor (*ST18*), apoptosis (*EMP1*), adherens junction (*CTNNA2*), calcium channel (*CACNB4*), and cytochrome P450 monooxygenase (*CYP11A1*) genes.

### Anti-Angiogenic VASH1 Mediated Signaling and Functional Enrichment in Angiogenesis Regulation

Since *VASH1* is an anti-angiogenic factor, and its gene expression displayed a positive correlation with BP, we next sought to investigate the expression levels of other candidate genes related to the *VASH1* pathway i.e, angiogenesis and tubulin polymerization [*AKT1*, *VEGF*, *HIF*-1, *TP53*, *SVBP*, *TTL*, *VASH2*, *TCP*, *SIRT1*, *CTGF*, *FGF2*]. Interestingly, except *VASH1*, none of these genes displayed a significant association with SBP or LVMI. Contrary to *VASH1,* which possesses anti-angiogenic and tubulin carboxypeptidase activity, *VASH2 is* pro-angiogenic, and *TTL* exhibits tubulin tyrosine ligase activity.

The significant positive enhancement of *VASH2* and *TTL* was only noted in the mid-BP group; the expression of these genes decreased sequentially in the high-BP and then the high-BP+high-LVMI group as expected. Conversely, *SVBP* levels increased sequentially when going from mid-BP to high-BP to high-BP+highLVMI (Fig. 3A). *FGF2* levels were diminished in increasing LVMI but failed to reach significance (data not shown). Functional gene enrichment using Metascape revealed that the pathways related to circulatory system processes, the vascular system, BP regulation, the TGF-β signaling pathway, and muscle contraction were enriched in upregulated gene sets (Fig. 3B, Supplemental File S3) and downregulated gene sets (Fig. 3C, Supplemental File S3). Thus, the transcriptomic analysis identified *VASH1/SVBP* as a critical factor with increased expression in response to BP that acts through its inhibition of angiogenesis signal and tubulin carboxypeptidase activity.

**Fig. 3.**
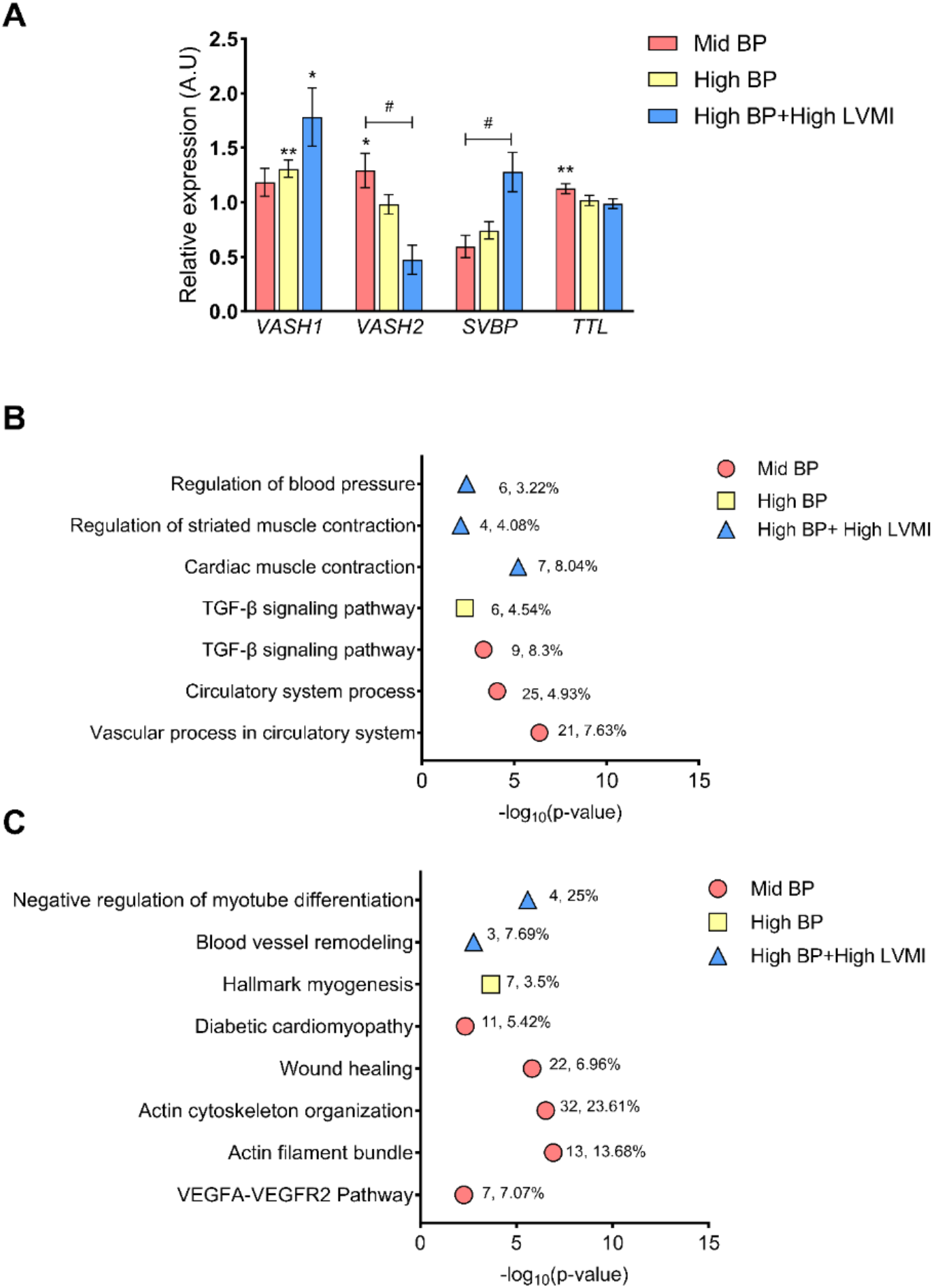
Expression pattern of the angiogenesis-related genes and their functional enrichment with increasing CVD risk phenotypes. (**A)** The fold change of expression for the candidate genes involved in angiogenesis in mid-BP, high-BP, high-BP+high-LVMI groups in comparison to their respective control groups (low-BP, low-BP, and low-BP+low LVMI respectively). The fold change is calculated using average FPKM values or normalized sequencing read counts in each group normalized to their average of respective control groups and plotted as Mean ± S.E.M. * represents p-value< 0.05, ** p-value <0.01 when comparing values using a two-tailed, unpaired students t-test. **B-C,** Summary of functional enrichment of the up-regulated **(B),** and down-regulated genes **(C)** to the angiogenesis and muscle cell-related pathways/categories. From the top 100 gene sets generated by Metascape, only those related to angiogenesis, cell adhesion, and muscle-cell-related pathways having a significant p-value of <0.01 (log 10 P value 0.01=-2) were visualised. The number of DEGs enriched for the pathway and its percentage enrichment are labelled against each enrichment category. A.U: Arbitrary Units, DEG: Differentially expressed genes; FPKM: Fragments Per Kilobase of transcript per Million mapped reads, LVMI: Left Ventricular Mass Index

### Small RNA-Mediated Gene Regulation in Circulating PBMCs

To study the epigenetic and post-transcriptional regulation of marker genes and associated signaling pathways, total miRNAs were utilized to perform Small RNA Sequencing. Small RNA Sequencing analysis followed by DESeq2 analysis revealed numerous differentially regulated miRNAs in the high-BP+high-LVMI (35 upregulated, 10 downregulated), mid-BP (13 upregulated, 14 downregulated), and high-BP (9 upregulated, 13 downregulated) groups as compared to the low BP+low LVMI and low BP groups (Fig. 4A-C, Supplemental File S1). Similar to our RNA Seq data (described above), miRNA expression differences also failed to reach statistical significance after FDR correction and were selected based on the significance threshold of P-value ≤ 0.05 and log_2_fold change cutoff 0.5. We next compared the dysregulated miRNAs with significance threshold of P-value ≤ 0.05 and without applying any log Fold change cutoff and observed that only 1 miRNA from each mid-BP group and high-BP group were correspondingly upregulated in the high-BP+high-LVMI group. There were no miRNAs that were commonly downregulated among mid-BP, high-BP, and high-BP+high-LVMI groups (Fig. 4D & E). These results suggest that miRNA signatures are distinct and unique to the BP and LVMI groups, suggesting their different roles in BP and CV-TOI.

**Fig. 4.**
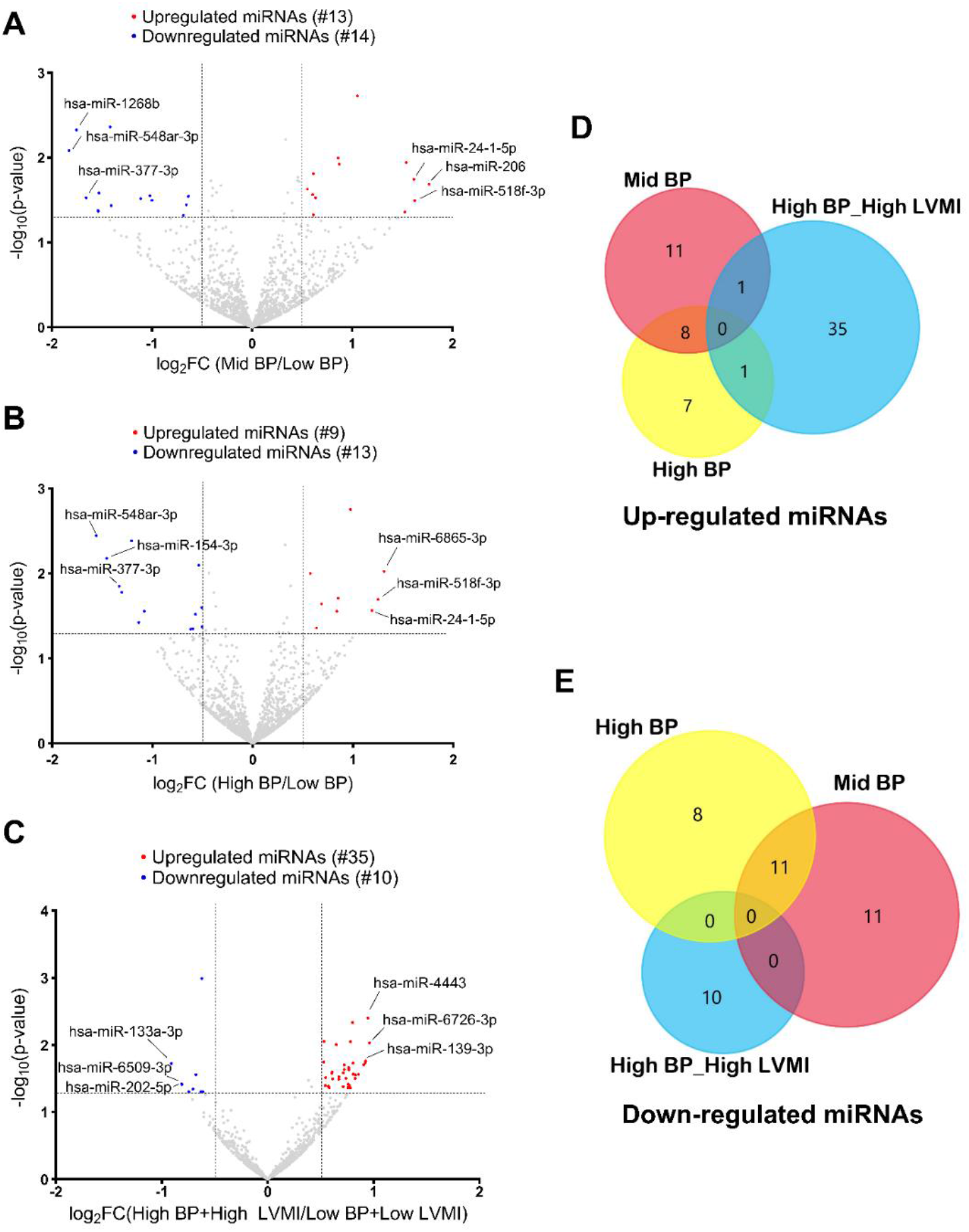
Micro RNA expression pattern in PBMCs from SHIP AHOY participants with mid-BP, high-BP, and high-BP+high-LVMI. Volcano Plot representing differentially expressed miRNAs in mid-BP group **(A)**, high-BP group **(B),** and high-BP+high-LVMI group **(C),** each compared to their respective controls. MicroRNAs with changes in expression and their p-value are determined by DEseq2 algorithm. The red color represents the increased miRNA expression, and the blue represents the diminished miRNA expression with p-value <0.05 and an absolute value of log_2_ fold change cut off 0.5. The top 3 up-regulated and down-regulated miRNAs were labeled for each analysis. The upregulated and downregulated miRNAs are compared among mid-BP, high-BP and high-BP+high-LVMI groups. **D-E,** Venn diagrams showing the upregulated set of miRNAs with a p-value<0.05 **(D),** and downregulated set of miRNAs with a p-value<0.05 (**E).** BP: Blood pressure; LVMI: Left ventricular mass index; miRNA: MicroRNAs; PBMCs: Peripheral blood mononuclear cells; SHIP AHOY = Study of High Blood Pressure in Pediatrics: Adult Hypertension Onset in Youth

### Prediction of miRNA Binding Sites in VASH1 3’UTR region

As our RNA sequence analysis revealed a strong association of *VASH1* gene upregulation with BP, we next investigated the association of miRNAs that regulate *VASH1* gene post-transcriptionally. TargetScan predicted potential binding sites for 153 miRNAs covering all conserved and poorly conserved sites (Supplemental File S4). As RNA and miRNA expressions are negatively correlated, we examined whether these miRNAs were among the downregulated miRNAs of mid-BP, high-BP, and high-BP+ high-LVMI groups (Fig. 5, Upper Panel). The miRNAs, miR-335-5p, miR-660-5p from high-BP+high-LVMI group and miR-377-3p, miR-760, miR-24-3p from the Mid-BP and high-BP group were significantly downregulated. Within these 5 miRNAs, the miR-24-3p binding site was a conserved site among vertebrates, miR-377-3p, miR-760, and miR-335-5p binding sites were conserved among mammals, whereas the miR-660-5p binding site was predicted to be a poorly conserved site (Fig. 5, Lower Panel). These findings suggest a potential post-transcriptional role for miR-24-3p in *VASH1* gene expression. Next, we used the miRNA expression values of the miRNAs from individual participants to identify if miRNA expression correlates with blood pressure and LVMI. Among 153 predicted *VASH1* miRNAs; miR-335-5p and miR-500a-3p displayed a significant negative correlation with blood pressure (Table 4). Similarly, miR-30d-5p, miR-30e-5p, miR-140-3p, miR-146a-5p, miR-338-3p, miR-10b-5p, miR-532-3p, miR-320a, miR-629-5p, miR-500a-3p and miR-942-5p were negatively correlated with LVMI (Table 5). However, except miR-335-5p none of the downregulated *VASH1* miRNAs identified using DESeq2 analysis failed to show any significant correlation with blood pressure or LVMI. Collectively, the downregulation of *VASH1* miRNAs correlates with the increased expression levels of *VASH1* mRNA in circulation. This suggests the miRNA mediated post-transcriptional regulation of *VASH1* in circulation may mediate anti-angiogenic pathways under conditions of elevated BP and its de regulation might contribute to CV-TOI.

**Fig. 5.**
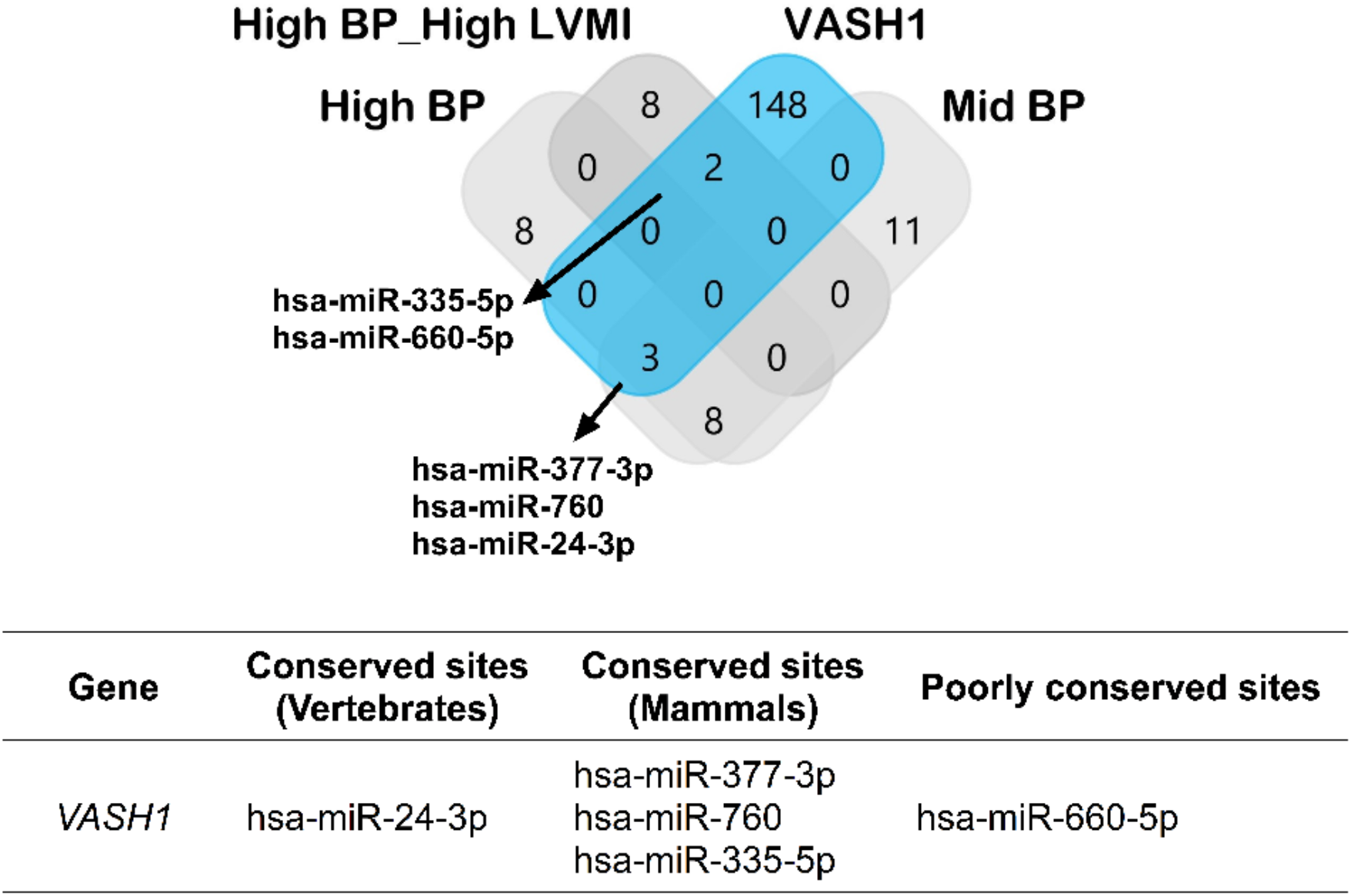
Potential miRNAs involved in post-transcriptional gene silencing of *VASH1*. ***<u>Upper panel</u>*:** The miRNAs for *VASH1* were predicted using Target Scan and compared with the down-regulated miRNA sets from mid-BP, high-BP and high-BP+high-LVMI groups. ***<u>Lower panel</u>:*** The predicted miRNAs were classified according to their binding site conservation based on the conservation cut-offs described by the target scan prediction tool.

**Table 4.** Correlation of predicted *VASH1* miRNAs with systolic blood pressure percentile.

| Predicted *VASH1*<br>miRNAs* (TPM) | Systolic BP percentile |  | n |
| --- | --- | --- | --- |
|  | Spearman r | p-value |  |
| hsa-miR-335-5p | -0.1891 | **0.0489** | 109 |
| hsa-miR-500a-3p | -0.2111 | **0.0332** | 102 |

\*Expression mean as determined from the miRNA seq data of peripheral blood mononuclear cells
TPM: transcripts per million

**Table 5.** Correlation of predicted *VASH1* miRNAs with left ventricular mass index.

| Predicted *VASH1*<br>miRNAs* (TPM) | LVMI (g/m<sup>2.7</sup>) |  | n |
| --- | --- | --- | --- |
|  | Spearman r | p-value |  |
| hsa-miR-30d-5p | -0.2162 | **0.024** | 109 |
| hsa-miR-30e-5p | -0.231 | **0.0156** | 109 |
| hsa-miR-140-3p | -0.2117 | **0.0271** | 109 |
| hsa-miR-146a-5p | -0.1902 | **0.0476** | 109 |
| hsa-miR-338-3p | -0.3226 | **0.0327** | 44 |
| hsa-miR-10b-5p | -0.2161 | **0.0254** | 107 |
| hsa-miR-532-3p | -0.2532 | **0.0079** | 109 |
| hsa-miR-320a | -0.2008 | **0.0363** | 109 |
| hsa-miR-629-5p | -0.2009 | **0.0362** | 109 |
| hsa-miR-500a-3p | -0.2155 | **0.0296** | 102 |
| hsa-miR-942-5p | -0.2464 | **0.0098** | 109 |

\*Expression mean as determined from the miRNA seq data of peripheral blood mononuclear cells
LVMI: left ventricular mass indexed to height; TPM: transcripts per million

### Whole Genome Methylation as an Epigenetic Regulator of BP and LVMI-associated Differential Expressed Genes

The major epigenetic mechanisms in complex disease pathogenesis include methylation, histone acetylation, and expression of non-coding RNAs including miRNAs. As highlighted in the previous section, potential miRNAs with predicted binding sites in *VASH1* were downregulated, suggesting a possible loss of the inhibitory effect of these miRNAs in *VASH1* mRNA levels. To further explore the epigenetic regulators of *VASH1,* DNA isolated from PBMCs of participants in the mid-BP and high-BP+high-LVMI groups were subjected to methylation sequencing and compared with low BP+low LVMI participants in a pair wise comparison. The differentially methylated regions (DMRs) were identified based on the number of CpG islands and the depth of the methylation using Fisher exact test. There were numerous differences among the sample comparisons in the number of DMR genes; to ensure robust and meaningful analysis, we focused on DMR genes that consistently showed hyper- or hypo- methylation across multiple comparisons. Specifically, we included only those DMRs that exhibited consistent methylation changes (hyper- or hypo-methylation) in at least 6 of the 10 pairwise comparisons. We excluded DMRs with equal hyper- and hypo-methylation patterns across the comparison groups to avoid ambiguous results. Among the selected DMRs, the total number of hypomethylated genes was higher in the mid-BP group in comparison to the high-BP+high-LVMI group, suggesting a significant enrichment of whole genome hypermethylation with increasing BP and LVMI (Fig. 6A, Supplemental File S5). Among these mid-BP DMR genes, *SH3BP4* displayed hypomethylation in all comparisons and *POTEI*, *SASH1, STK32C* and *CYP2E1* exhibited hypermethylation in at least 8 out of 10 comparisons (Fig. 6B, Supplemental File S5). Similarly, in the high-BP+high-LVMI group, the DMRs *ADARB2*, *MADIL1, and PRDM16 displayed hypomethylation, and GSE1, RUNX1, MN1,* and *VAV2* had hypermethylation patterns (Fig. 6C, Supplemental File S5).

**Fig. 6.**
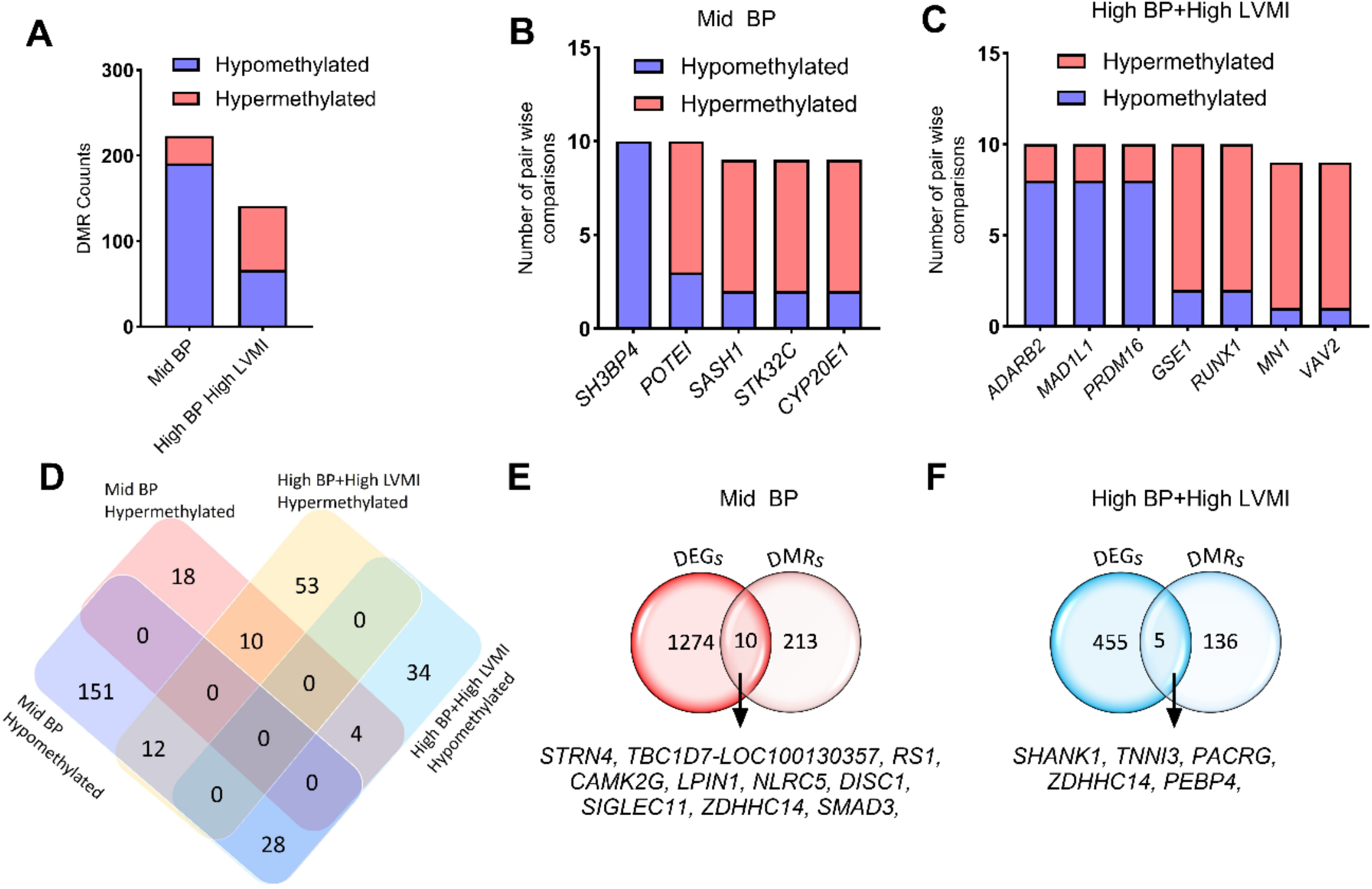
Methylation pattern of whole genome DMRs and relation with RNA expression levels in SHIP AHOY participants with mid-BP and high-BP+high-LVMI. **A,** Bar graph displaying the number of hypermethylated and hypomethylated DMR genes ranked based on their appearance in at least 6 pairwise comparisons in mid-BP (n=10), and high-BP+high-LVMI groups (n=10) each compared to low-BP+low-LVMI (n=10). **B-C,** Bar graph showing the hypermethylated and hypomethylated DMR genes that appeared in the maximum number of comparisons in the **(B)** mid-BP (n=10), and **(C)** high-BP+high-LVMI groups (n=10). **D,** Venn diagram showing the comparison of significant hyper- and hypomethylated DMRs between the mid-BP and high-BP+high-LVMI groups. **E-F,** Venn diagram comparing the DEGs and DMRS in mid-BP **(E)**, and high-BP groups **(F)**. DMRs are selected based on their significant difference in the All CpG average methylation rate of mid-BP and high-BP+high-LVMI samples in comparison to their low-BP+low LVMI samples with a p-value<0.05 (Fisher exact test). The DMRs are then ranked based on their appearance in the 10 pairwise comparisons; only those having a minimum of 6 comparisons were considered for further analysis. BP: blood pressure; DEG: differentially expressed genes; DMR: differentially methylated region; LVMI: left ventricular mass index; SHIP AHOY: Study of High Blood Pressure in Pediatrics: Adult Hypertension Onset in Youth

We also analyzed DMRs between the mid-BP and high-BP+high-LVMI groups to identify potential hypo- or hypermethylated genes in either or both groups. This revealed that 10 DMR genes were hypermethylated and 28 DMR genes were hypomethylated in both mid-BP and high-BP+high-LVMI. Whereas 12 DMR genes that were hypomethylated in mid-BP were hypermethylated in high-BP+high-LVMI, and 4 genes hypermethylated in mid-BP were hypomethylated in high-BP+high-LVMI (Fig. 6D, Table 6). Interestingly genes that demonstrated DMR in the maximum number of pair-wise comparisons (Fig. 6B-C) viz., *GSE1*, *POTE1*, *MN1*, *MAD1L1*, *SH3BP4*, *RUNX1*, *VAV2*, *ADARB2,* and *STK32C* displayed hypo- or hypermethylation in both groups or vice versa (Table 6). Functional enrichment of the DMRs among the mid-BP and high-BP+high-LVMI groups revealed that the genes related to blood pressure, cardiac function, and angiogenesis were significantly enriched (Table 7, Supplemental File S6). Interestingly, the DMR genes of the mid-BP group showed enrichment for “process related to cardiac muscle,” inferring that the CV-TOI process was initiated at the early stage of BP rise.

**Table 6.** Genes with significant differentially methylated regions in adolescents with increased left ventricular mass index and BP >80^th^ percentile.

| Category | List of genes |
| --- | --- |
| Hypermethylated in mid-BP and high-BP+high-LVMI | **GSE1**, **POTE1**, ALK, ZDHHC14, TSHZ2, LITAF, RPS6KA2, OPCML, **MN1**, GLT1D1 |
| Hypomethylated in mid-BP and high-BP+high-LVMI | PDHA1, SEPTN6, ARHGAP6, STARD8, CLCN4, ADAP1, ALG1L1P, SNX29, XPNPEP2, AATK, NHSL1, DPEP1, STK10, NFATC1, JARID2, RPTOR, PBX1, FOXP1, **MAD1L1**, SLC45A4, SORCS2, PRKAG2, KLHL29, AMZ1, RS1, RIN3, **SH3BP4**, NLRC5 |
| Hypomethylated in mid-BP and Hypermethylated in high-BP+high-LVMI | **RUNX1**, GNG7, **VAV2**, RXRA, TRAPPC9, LRP5, ANKRD33B, BCR, RAI1, RAD51B, COTL1, DOCK1 |
| Hypermethylated in mid-BP and Hypomethylated in high-BP+high-LVMI | PTPRN2, **ADARB2**, **STK32C**, CSMD1 |

Genes highlighted in bold are significant DMRs that appeared in the maximum number of comparisons either in mid-BP vs low-BP + low-LVMI or high-BP + high-LVMI vs low-BP + low-LVMI

**Table 7.** Functional enrichment of genes with significant differentially methylated regions (DMRs) among mid-BP and high-BP+high-LVMI group SHIP AHOY participants.

| Category | Term | Description | LogP | Log(q-value) | Percentage of DMR genes enriched |
| --- | --- | --- | --- | --- | --- |
| **Mid-BP DMRs** |  |  |  |  |  |
| GO Biological Processes | GO:0090257 | Regulation of muscle system process | -5.68 | -2.55 | 4.49 |
| GO Biological Processes | GO:0030029 | Actin filament-based process | -5.53 | -2.51 | 2.78 |
| GO Biological Processes | GO:0008015 | Blood circulation | -4.87 | -2.10 | 3.12 |
| WikiPathways | WP3888 | VEGFA VEGFR2 signaling | -3.44 | -1.19 | 2.55 |
| GO Biological Processes | GO:0010595 | Positive regulation of endothelial cell migration | -3.38 | -1.14 | 4.58 |
| GO Biological Processes | GO:0008217 | Regulation of blood pressure | -3.34 | -1.13 | 3.76 |
| GO Biological Processes | GO:0045766 | Positive regulation of angiogenesis | -3.33 | -1.12 | 3.74 |
| GO Biological Processes | GO:0098901 | Regulation of cardiac muscle cell action potential | -2.92 | -0.86 | 10.35 |
| GO Biological Processes | GO:0097435 | Supramolecular fiber organization | -2.86 | -0.83 | 2.04 |
| GO Biological Processes | GO:0014902 | Myotube differentiation | -2.63 | -0.67 | 5.26 |
| GO Biological Processes | GO:0055117 | Regulation of cardiac muscle contraction | -2.61 | -0.67 | 5.19 |
| Reactome Gene Sets | R-HSA-5576891 | Cardiac conduction | -2.54 | -0.62 | 3.79 |
| GO Biological Processes | GO:1903522 | Regulation of blood circulation | -2.53 | -0.61 | 2.71 |
| GO Biological Processes | GO:0086091 | Regulation of heart rate by cardiac conduction | -2.51 | -0.60 | 7.5 |
| GO Biological Processes | GO:1903115 | Regulation of actin filament-based movement | -2.51 | -0.60 | 7.5 |
| KEGG Pathway | hsa05412 | Arrhythmogenic right ventricular cardiomyopathy | -2.47 | -0.57 | 4.76 |
| Reactome Gene Sets | R-HSA-397014 | Muscle contraction | -2.39 | -0.52 | 2.93 |
| KEGG Pathway | hsa05410 | Hypertrophic cardiomyopathy | -2.25 | -0.44 | 4.12 |
| Reactome Gene Sets | R-HSA-4420097 | VEGFA-VEGFR2 Pathway | -2.22 | -0.42 | 4.04 |
| GO Biological Processes | GO:0002027 | Regulation of heart rate | -2.14 | -0.39 | 3.85 |
| **High-BP+high-LVMI DMRs** |  |  |  |  |  |
| GO Biological Processes | GO:0003073 | regulation of systemic arterial blood pressure | -4.08 | -1.13 | 5.26 |
| GO Biological Processes | GO:0008217 | regulation of blood pressure | -3.61 | -0.89 | 3.23 |
| GO Biological Processes | GO:0003013 | circulatory system process | -3.19 | -0.66 | 1.78 |
| GO Biological Processes | GO:0055021 | regulation of cardiac muscle tissue growth | -2.42 | -0.29 | 4.48 |

As we did not observe *VASH1* or *VASH1* pathway gene expression to be enriched in the DMRs, we then compared the DEGs with the DMR genes to determine if any were epigenetically regulated. Our analysis showed that 10 DEGs in the mid-BP group (Fig. 6E, Supplemental File S5) and 5 DEGs in the high-BP+ high-LVMI (Fig. 6F) group might be regulated through their differential methylation pattern. Taken together, our global DNA methylation analysis of PBMC DNAs from mid-BP and high-BP+high-LVMI groups indicated that there is increasing hypermethylation of genes in the high-BP+high-LVMI group and identified candidate DMR genes associated with blood pressure, cardiac function, and angiogenesis. This also revealed that *VASH1* and *VASH1* pathway genes are likely not regulated through DNA methylation and showed distinct DMR profiles for mid-BP and high-BP+high-LVMI groups.

### Circulatory Proteins in the Pathogenesis of CV-TOI

Serum samples of SHIP AHOY participants were isolated, and protein profiling via mass spectrometry identified 298 circulating proteins (Supplemental File S7), with analyses that revealed several differentially regulated proteins in the mid-BP (Fig. 7A, Table 8) and high-BP+high-LVMI group as compared to the low BP+low LVMI group (Fig. 7B, Table 8). We found 22 circulatory proteins with a differential expression in the mid-BP and high-BP+high-LVMI groups with a significance threshold of P-value ≤ 0.05 and without applying any log Fold change cutoff. Among these circulatory proteins, 4 DEPs such as SOD3, IGFALS, PF4V1, and PROZ were similarly upregulated or downregulated in both the mid-BP and high-BP+high-LVMI groups (Fig. 7C) and displayed a significant differential expression either in both the groups or in high-BP+high-LVMI group (Fig. 7D). SOD3 and IGFALS levels were diminished and PF4V1 and PROZ levels were elevated in the mid-BP and high-BP+high-LVMI in comparison to low BP+low LVMI group (Fig. 7D). A fold change cut off>0.5 yielded 4 (APOL1, H1-3, GSN and PROZ) significantly upregulated proteins and 6 downregulated proteins in the mid-BP group (Fig. 7A, Table 8). Similarly, 7 proteins (PROZ, PF4, BTD, PRG4, CTSA, APOC3, MAN1A1) were upregulated, and 5 proteins were downregulated (JCHAIN, SOD3, FGG, CD5L, KRT10) in the high-BP+high-LVMI group (Fig. 7B, Table 8). Functional enrichment of these DEPs revealed various associated pathways related to inflammation, immune response, Insulin growth factor, oxidative stress and extracellular matrix-related pathways and processes (Table 9, Supplemental File S8), among which the DEPs of high-BP+high-LVMI group are known to contribute more towards inflammatory response.

**Fig. 7.**
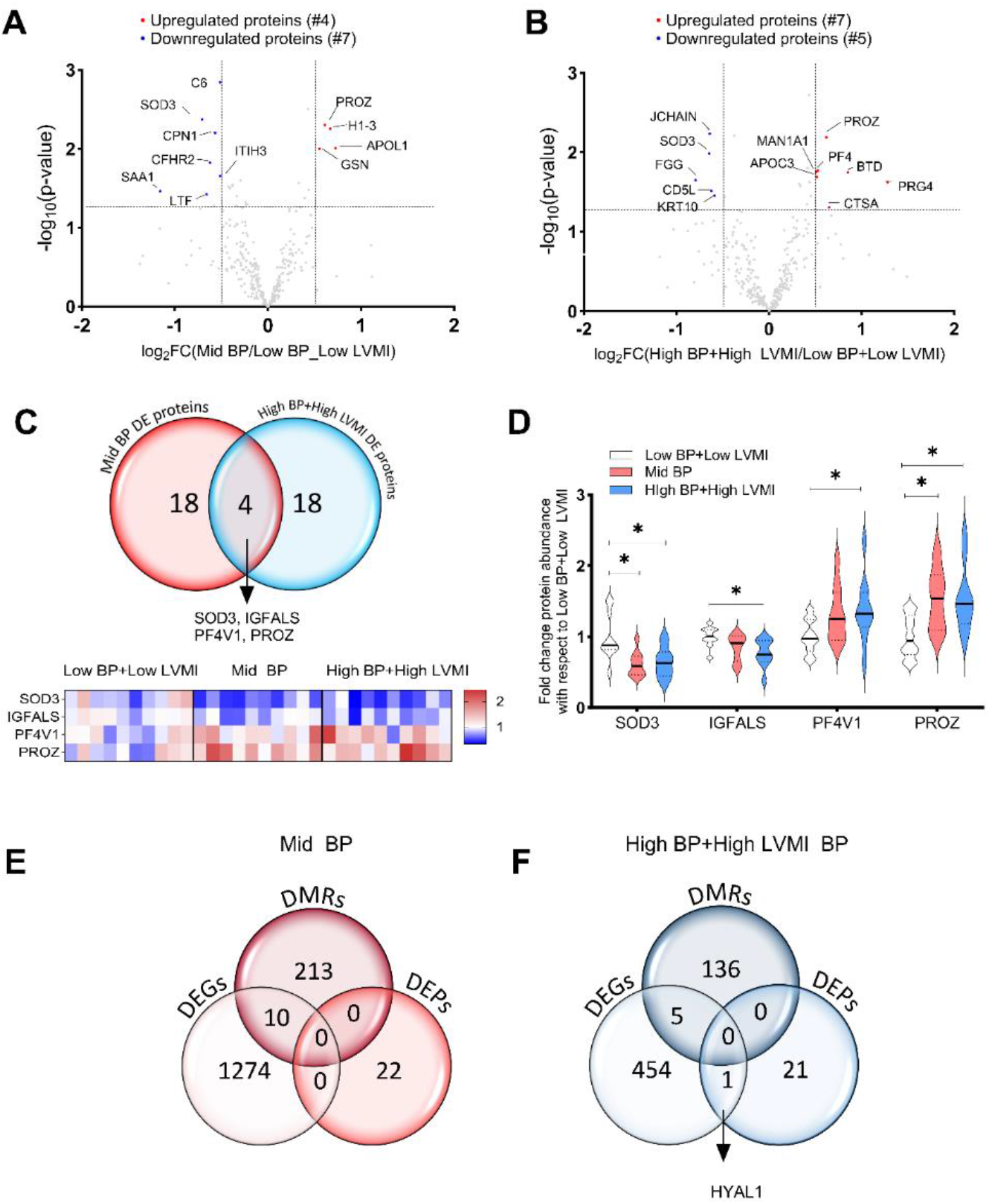
Differentially expressed proteins (DEPs) in the serum of SHIP AHOY participants with mid-BP and high-BP+high-LVMI. Volcano Plot representing differentially expressed proteins with a log _2_ fold change cut-off 0.5 in mid-BP **(A)** and, high-BP+high-LVMI groups **(B)**. The log_2_ fold change of average protein abundance in mid-BP (n=10), and high-BP+high-LVMI groups (n=10) in comparison to low-BP+low LVMI (n=10) were plotted against -log_10_ p-value. The red color represents the significantly up-regulated proteins, whereas the blue represents the down-regulated proteins. **C,** Venn diagram representing DEPs in mid-BP and high-BP+high-LVMI group identifies SOD3, IGFALS, PF4V1 and PROZ as common DEPs under high blood pressure conditions. **D,** Violin plots showing the expression levels of SOD3, IGFALS, PF4V1 and PROZ proteins in mid-BP and high-BP+high-LVMI groups as mean ± S.E.M. * representing p-value<0.05, of unpaired, two-tailed multi variable t-test. **E-F,** Venn diagram showing the DEGs, DMRs, and DEPs in mid-BP **(E)** and high-BP+high-LVMI groups of SHIP AHOY study participants **(F)**. BP: blood pressure; DEG: differentially expressed genes; DMR: differentially methylated regions; LVMI: left ventricular mass index; SHIP AHOY: Study of High Blood Pressure in Pediatrics: Adult Hypertension Onset in Youth

**Fig. 8.**
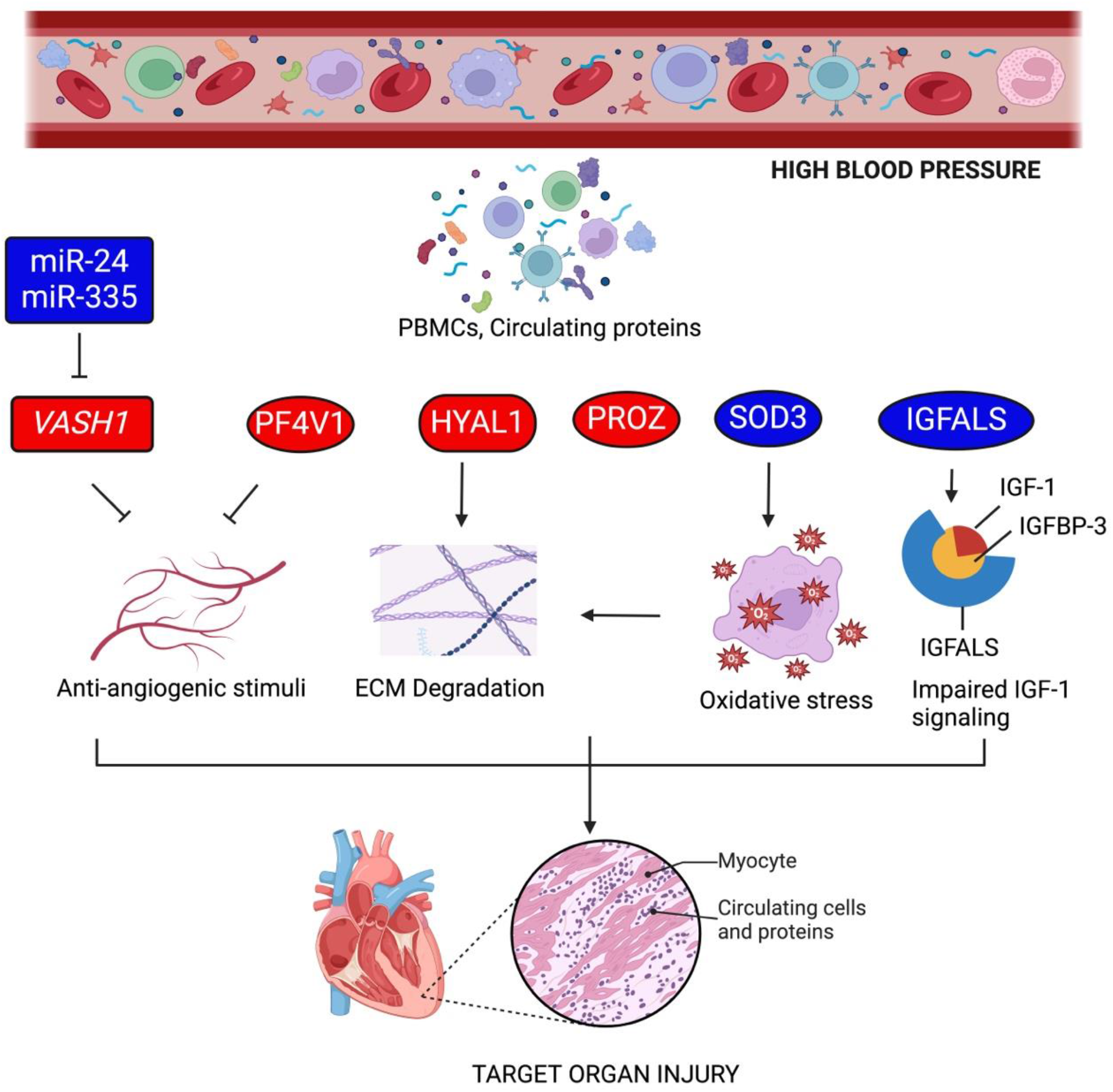
Proposed mechanism of target organ injury in youth with high blood pressure. Based on the RNA, miRNA, and methylation sequencing and proteomic analysis of mid-BP, high-BP and high-BP+high-LVMI youth subjects, anti-angiogenic stimuli and ECM degradation through the signals from PBMCs and circulating proteins leads to target organ injury. Rectangular boxes are the genes and miRNAs displaying differential transcript level in PBMCs; oval boxes represent differential protein levels in circulation; cylindrical boxes represent genes differentially expressed in PBMCs and proteins differentially expression in circulation. Red color represents up-regulation and blue color indicates down-regulation (Figure Created with BioRender.com).

**Table 8.** List of differentially expressed proteins in adolescents with increased left ventricular mass index and BP >80^th^ percentile.

| Mid-BP Vs low-BP+low-LVMI |  |  | High-BP+high-LVMI Vs low-BP+low-LVMI |  |  |
| --- | --- | --- | --- | --- | --- |
| Gene Symbol | log2 FC | -log10 (P-value) | Gene Symbol | log2 FC | -log10 (P-value) |
| SAA1 | -1.16 | 1.46 | FGG | -0.79 | 1.65 |
| SOD3 | -0.71 | 2.37 | SOD3 | -0.64 | 1.99 |
| LTF | -0.66 | 1.42 | JCHAIN | -0.64 | 2.24 |
| CFHR2 | -0.62 | 1.83 | CD5L | -0.62 | 1.52 |
| CPN1 | -0.57 | 2.21 | KRT10 | -0.59 | 1.45 |
| ITIH3 | -0.52 | 1.66 | A2M | -0.48 | 1.44 |
| C6 | -0.52 | 2.84 | IGFALS | -0.38 | 2.21 |
| HSPA5 | -0.43 | 1.52 | HYAL1 | 0.33 | 1.60 |
| CALR | -0.41 | 1.50 | IGFBP3 | 0.34 | 1.57 |
| SPP2 | -0.38 | 1.53 | KLKB1 | 0.40 | 2.12 |
| C4B | -0.38 | 1.55 | RARRES2 | 0.41 | 1.34 |
| OIT3 | -0.34 | 1.40 | PPBP | 0.42 | 1.80 |
| C1R | -0.30 | 1.60 | CPB2 | 0.43 | 2.72 |
| IGFALS | -0.24 | 1.33 | APOA4 | 0.44 | 1.63 |
| QSOX1 | 0.33 | 1.57 | PF4V1 | 0.45 | 1.45 |
| PF4V1 | 0.40 | 1.38 | MAN1A1 | 0.51 | 1.75 |
| SERPINA1 | 0.43 | 2.50 | APOC3 | 0.52 | 1.69 |
| FBLN5 | 0.44 | 1.48 | PF4 | 0.53 | 1.77 |
| GSN | 0.55 | 1.99 | PROZ | 0.62 | 2.19 |
| PROZ | 0.61 | 2.30 | CTSA | 0.65 | 1.31 |
| H1-3 | 0.67 | 2.25 | BTD | 0.86 | 1.74 |
| APOL1 | 0.73 | 2.01 | PRG4 | 1.28 | 1.63 |

**Table 9.** Functional enrichment of significantly dysregulated serum proteins (Differentially expressed proteins; DEPs) among mid-BP and high-BP+high-LVMI group SHIP AHOY participants.

| Category | Term | Description | LogP | Log(q-value) | Percentage of DEP genes enriched |
| --- | --- | --- | --- | --- | --- |
| **Mid-BP DEPs** |  |  |  |  |  |
| Reactome Gene Sets | R-HSA-977606 | Regulation of Complement cascade | -9.73 | -5.47 | 10.64 |
| Reactome Gene Sets | R-HSA-381426 | Regulation of Insulin-like Growth Factor (IGF) transport and uptake by Insulin-like Growth Factor Binding Proteins (IGFBPs) | -9.51 | -5.47 | 5 |
| GO Biological Processes | GO:0042060 | Wound healing | -2.83 | -0.20 | 0.95 |
| **High-BP+high-LVMI DEPs** |  |  |  |  |  |
| GO Biological Processes | GO:0006954 | Inflammatory response | -8.47 | -4.43 | 1.42 |
| GO Biological Processes | GO:0009611 | Response to wounding | -6.36 | -3.13 | 1.42 |
| GO Biological Processes | GO:0042060 | Wound healing | -5.57 | -2.45 | 1.58 |
| GO Biological Processes | GO:1990266 | Neutrophil migration | -4.40 | -1.61 | 3.26 |
| GO Biological Processes | GO:0097530 | Granulocyte migration | -4.29 | -1.55 | 3 |
| Reactome Gene Sets | R-HSA-381426 | Regulation of Insulin-like Growth Factor (IGF) transport and uptake by Insulin-like Growth Factor Binding Proteins (IGFBPs) | -4.00 | -1.27 | 2.4 |
| WikiPathways | WP5095 | Overview of proinflammatory and profibrotic mediators | -3.96 | -1.26 | 2.33 |
| GO Biological Processes | GO:0000302 | Response to reactive oxygen species | -3.54 | -0.93 | 1.68 |
| Reactome Gene Sets | R-HSA-1474244 | Extracellular matrix organization | -2.89 | -0.38 | 1 |
| GO Biological Processes | GO:0006979 | Response to oxidative stress | -2.65 | -0.18 | 0.83 |

Correlation and gene cluster analyses of DEGs, DMRs and DEPs of mid-BP and high-BP+high-LVMI groups indicated that they are distinct except for *HYAL1*. *HYAL1* transcript levels and protein levels were upregulated in circulating PBMCs and the serum of the high-BP+high-LVMI group (Fig. 7E-7F). Overall, these data suggest dysregulation in serum anti-angiogenic factor (PF4V1), insulin pathway (IGFALS), oxidative stress (SOD3), and ECM (HYAL1) pathways likely account for the induction of pathologic CV remodeling in participants with high-BP and CV-TOI.

## DISCUSSION

In this observational study, we evaluated circulatory markers and associated signaling pathways putatively associated with CV-TOI to explore the potential origins of sequelae related to high-BP in youth. Specifically, to understand the regulation of circulatory factors in youth with high-BP and high-LVMI, a form of pathological cardiac remodeling, PBMCs, and serum samples from youth enrolled in the SHIP AHOY study underwent molecular and pathway analysis via RNA-Seq, miRNA-Seq, global DNA methylation, and mass spectrometry. RNA Seq analyses revealed a possible positive correlation between circulatory antiangiogenic factor vasohibin-1 (*VASH1*), blood pressure and cardiac remodeling. Further, these findings and other analyses highlight that *VASH1* is not only likely a key player in associated pathological processes leading to the development of LVH in patients but that other potential targets such as antiangiogenic stimuli through PFV1, ECM degradation through HYAL1, antioxidative pathways through SOD3, and impaired insulin signaling pathway in circulation may each facilitate remodeling of cardiomyocytes, potentially impacting their contractility.

### Anti-angiogenesis: A Key Player in Hypertension-Induced TOI through VASH1

While the current study detected the involvement of several circulatory master genes and pathways in young patients with high-BP and CV-TOI characterized by cardiac hypertrophy, the antiangiogenic factor *VASH1* stood out as one of the top differentially regulated genes, mainly due to its positive correlation with SBP (Fig. 2C) and elevated expression level in high-BP (Fig. 3A) and high-BP+high-LVMI group (Fig. 3A). *VASH1* has been related to hypertension^21^ and hypertensive cardiac remodeling in animal and adult human studies.^22^ *VASH1* is considered to be part of the vascular “self-defense” system by promoting endothelial cell stress tolerance and preventing age-related vascular disease manifesting as increased arterial stiffness early in the clinical course of CV disease. In response to cellular stress, as seen with hypertension, endothelial cells increase VASH1 gene expression.^23^ The link between the endothelial and cardiomyocyte actions of *VASH1* may be explained by findings reported in heart transplant recipients: heart transplant patients with pathologic cardiac remodeling exhibit a significant decrease in the expression and capillary density of VEGF-A and vascular endothelial growth factor receptor 1 (VEGFR1),^24^ suggesting impaired cardiac angiogenesis.

Supporting this hypothesis, recent studies by Chen et al. have revealed the functional restorative effect of VASH1 genetic inhibition in failing human CMs,^25^ suggesting a noteworthy involvement of VASH1 in cardiac dysfunction and associated signaling. Corroborating these reports, our functional enrichment of DEGs (Fig. 3B and 3C) display a progressive development of TOI through upregulation in the vascular process (mid-BP), circulation process (mid-BP) >TGF-β Signaling (mid-BP and high-BP) > blood pressure and muscle contraction (high-BP+high-LVMI). On the other hand, downregulation of the VEGF pathway, actin structural protein, wound healing, myopathy-related genes (mid-BP)<Myogenesis (high-BP) <Blood vessel remodeling and negative regulation of myotube differentiation (high-BP+high-LVMI) in circulation accounts for the CV-TOI.

### Differential Expression of miRNAs as Potential Post-transcriptional Regulators of VASH1

To explore VASH1 regulation in detail, we examined the levels of different VASH1-targeted miRs in PBMCs. Our small RNA seq data also suggests a possible post-transcriptional regulation of VASH1 through microRNAs. RNA-Seq analyses from circulatory PBMCs revealed a significant upregulation in the antiangiogenic VASH1 gene (Fig. 2A and 3A), whereas *VASH1* predicted miRNAs viz., miR-24-3p, miR-335-5p, miR-660-5p, miR-377-3p and miR-760 display downregulation (Fig. 5). Our correlation analysis of TPM counts of these 5 miRNAs with blood pressure or LVMI did not achieve any significant correlation for miR-24-3p. Interestingly, miR-335-5p, displayed a negative correlation with blood pressure (Table 4). Based on the conservation score and correlation, miR-24-3p and miR-335-5p are expected to exhibit a stronger silencing action on *VASH1* gene expression. Contrary to the data published on the upregulation of miR-335,^26,27^ in hypertension and right ventricular remodeling and elevated levels of miR-24-3p levels in obese children with metabolic syndrome, positively correlated with various metabolic syndrome clinical parameters, including blood pressure.^28^ further validation is warranted to define its role in adult-onset CV-TOI. However, the expression level of miR-24-3p in PBMCs of CAD patients aged ∼60 years displayed downregulation, signifying its crucial role in the progression of atherosclerotic plaques.^29^ MicroRNA-4530 previously validated to target VASH1 and promote angiogenesis in breast cancer carcinoma cells,^30^ did not display any differential expression pattern in our small RNA sequencing. Thus, the downregulation of miR-24-3p and miR-335-5p highlights their role in the pathophysiology of CV-TOI. Further studies are necessary to validate the precise gene-silencing mechanism of *VASH1* by these miRNAs.

### Candidate VASH1 microRNAs as Predictors of Hypertension and Hypertension-induced CV-TOI

Our correlation of expression levels of in silico-predicted *VASH1* miRNAs with blood pressure and LVMI revealed a distinct set of miRNAs displaying a negative correlation with blood pressure and LVMI (Table 4 & 5). This suggests that their deregulation might affect the VASH1 transcript levels with respect to the changes in blood pressure and LVMI. Intriguingly, evidence support the significance of miR-378 and miR-361 in essential hypertension.^31–33^ Among the miRNAs with expression levels negatively correlated with LVMI, miR-30, miR-10, have previously been correlated with various cardiac-related pathological conditions.^34–37^ Thus, collectively, our RNA and small RNA seq data provide evidence for differential expression of anti-angiogenic gene *VASH1* and its predicted miRNAs. The differentially expressed miRNAs observed in our analysis are reportedly implicated as predictors of blood pressure and cardiac-associated disease phenotypes, particularly associated with left ventricular hypertrophy.

### Whole genome Methylation in Pathogenesis of Hypertension-induced CV-TOI

We also asked whether the deregulation of *VASH1* and its predicted miRNAs observed in our sequence analysis might be at least partially due to differential methylation. To support this our whole genome methylation analysis suggested that there was an increase in the proportion of genes hypermethylated in the high-BP+high-LVMI group. DNA methylation at the candidate gene locus has been linked to the progression of hypertension^38^ and hypertrophy,^39^ supporting differences we observed in the hyper and hypomethylated DMR genes. In the context of the angiogenesis and CV-TOI, among the DMR genes, key genes are *GSE1*, for which differential methylation has been previously reported in patients with left ventricular hypertrophy and atrial fibrillation,^40,41^ and *VAV2,* a guanine nucleotide exchange factor for Ras-related GTPases and modulate receptor-mediated angiogenic responses.

Though there is no evidence for *VAV2* methylation in blood pressure or cardiac diseases, it has been significantly associated with left ventricular hypertrophy, cardiac fibrosis, and hypertension.^42^ The other potential genes associated with DMRs in our study included *SH3BP*, *CYPE21*, *PRDM16*, and *RUNX1,* as either their methylation or their functions were previously known to be involved in cardiometabolic phenotypes.^43–45^ The functional enrichment reveals that angiogenesis, VEGF, blood pressure, and muscle-related pathway genes are differentially methylated. Further, we investigated if methylation acts as a potential epigenetic regulator of the *VASH1* gene expression. Our study revealed that *VASH1* does not possess any CpG islands with differential methylation, however, we discovered other potential DMR genes that may be involved in the blood pressure and CV-TOI process. Additional experiments and functional validation of the identified DMRs are needed to confirm our findings. Identification of cell-free DNA methylation might help us pinpoint the potential biomarker for cardiac injury,^46^ and this serves as a non-invasive way of detecting human cardiomyocyte death.^47^

### Circulating anti-angiogenic Factors in Hypertension-induced CV-TOI

In order to evaluate if the *VASH1* up-regulation in PBMC RNA levels affected the VASH1 protein levels in circulation, the proteomic profile of the serum samples was determined using the LCMS approach.

The protein profiling revealed that VASH1 protein levels were not differentially expressed. However, our analysis capitulated differential levels for PROZ, SOD3, PFV1, and IGFALS proteins in mid-BP and high-BP+ high-LVMI. Inconsistent reports for PROZ levels and its gene variants in pathogenesis of cardiovascular disease.^48–50^ It promotes angiogenesis under ischemic conditions (negative correlation (low levels) with pre-diabetes and diabetes conditions.^51^ It has angiogenic potential in ischemic conditions.^52^ However, its potential as a CV-TOI is uncertain due to the existence of different circulating levels with respect to age and gender.^53^ Platelet factor 4 (PF4, CXCL4) is a small chemokine protein released by activated platelets with major physiological functions to promote blood coagulation, but its antiangiogenic properties are proven.^54^ PF4 and PF4V1 candidature in blood pressure and CVD are well documented.^55,56^

Deficiency or reduced expression of SOD3, which is an extracellular antioxidant enzyme localized mainly to the extracellular matrix, has been implicated in blood pressure and cardio-related disease phenotypes; conversely, increased expression of SOD3 has been reported as protective.^57–59^ Further, SOD3 inhibition has been known to inhibit angiogenesis^60,61^ and have a role in immune response^62^ and hyaluronic homeostasis.^63^ Interestingly, in our study, we observed that HYAL1 RNA levels and its protein levels were elevated in the circulation of the high-BP+high-LVMI group. Hyal1 deficiency has been shown to have protective effects, and Hyal1 inhibition was suggested as a potential therapeutic target in CV disease.^64^ Higher circulating HA levels and elevated HYAL1 expression were consistently reported as correlated with cardio-metabolic phenotypes.^65,66^ Further, hyaluronan affects the force generation of cardiomyocytes, and hyaluronic metabolism is crucial for ECM integrity.^67^ Importantly, high molecular weight HA inhibits angiogenesis, and the HA degradation products induce angiogenesis,^68–70^ suggesting the presence of a complex regulatory role of HA in cardiovascular health and disease. Further, functional enrichment of DEPs may imply that pathways or processes related to inflammation, oxidative stress, wound healing, insulin signaling, extracellular matrix, and immune response are deregulated in hypertensive youth.

### Limitations of the Study

Our study has certain limitations. While multi-omics analysis of changes in molecular markers in youth with hypertension offers the advantage of identifying the cues of CV-TOI process, whether the expression differences observed in these peripheral samples mediate effects within the target organ is unclear. Further, the serum samples obtained have large heterogeneity, and the cellular composition of the blood is highly dynamic and confounding. Additionally, though our transcriptomic and proteomic data had statistically significant results for differentially expressed genes and proteins, theses did not reach significance after applying multiple hypothesis testing, which is commonly observed in many transcriptomic analyses of human subject data where they have a minimal number of DEGs after FDR.^71,72^ A limitation of our study is that the cellular composition of the blood is highly dynamic and acts as a confounding variable when using PBMCs and circulatory proteins to assess circulatory epigenetic/protein biomarkers. Thus, further studies are needed to validate our observations. Though the present study in hypertensive youth sought to define basic aspects of disease progression at an early age, dynamic temporal changes as CV-TOI progresses are essential to understanding its evolution completely.

## Conclusions

Our data suggest that PH, in the presence of common intermediate phenotypes, may lead to CV-TOI in hypertensive youth through impaired angiogenesis mediated by circulatory *VASH1*, its miRNAs, and associated signaling pathways. Additionally, the deregulation of angiogenesis due to differential methylation of *VAV2*, elevated PF4V1 and SOD3 protein levels, and extracellular matrix degradation by HYAL1 might contribute to the pathological remodeling of cardiac tissues. This study provides initial evidence of global RNA and protein profiles in the circulation associated with hypertension in youth and identifies the likely involvement and deregulation of canonical pathways related to angiogenesis, inflammation, oxidative stress, and extracellular matrix organization. These findings suggest that such pathways may ultimately lead to cardiovascular dysfunction. Further research is warranted to utilize these multi-omic findings to develop therapeutic interventions for CV-TOI aimed at promoting angiogenesis and protecting against extracellular matrix degradation in patients with PH.

## MATERIALS AND METHODS

### Study Population and Design

The design and methods of the SHIP AHOY (Study of High Blood Pressure in Pediatrics, Adult Hypertension Onset in Youth) study have been previously described.^10^ Briefly, 397 youth aged 11 to less than 19 years with BP levels across the BP spectrum were enrolled between 2015 to 2017 from five pediatric specialty clinics at academic medical centers (nephrology clinics at University of Texas Health Houston, Children’s Hospital of Philadelphia, University of Rochester Medical Center, and Seattle Children’s Hospital; one joint nephrology and cardiology clinic at Cincinnati Children’s Hospital Medical Center). Efforts were made to achieve body mass index (BMI) balance across BP groups by periodically checking differences in BMI percentile among groups and actively recruiting normotensive obese controls. Clinical data collected include demographics, anthropometrics, resting and ambulatory BP measured according to pediatric guidelines; 2-D guided M-mode for left ventricular mass index (LVMI), diastolic function and systolic strain measurements; and tonometer-based assessment of pulse wave velocity (PWV), a measure of central arterial stiffness.^10^ All participants provided blood samples for analysis. Approval for studies on human samples was obtained from the Institutional Ethical Review Board of Cincinnati Children’s Hospital Medical Center, the central IRB of record, and from the University of Rochester, New York, due to their inability to participate in the central IRB. Informed consent, including the use of blood samples for research, was obtained from participants 18 years of age and older or from parents/guardians along with participant assent for participants < 18 years of age according to local investigational review board requirements.

### SHIP AHOY Study Participants

The Systolic Hypertension In Pediatrics – Adult Hypertension Onset in Youth (SHIP-AHOY) study was a multi-center study of the effects of elevated BP in youth on target organ injury that recruited participants across the spectrum of BP from low to high-BP (N=397) as previously described.^10^ In the epigenetic sub-study of the SHIP AHOY study, participants were ranked based on their BP and LVMI. A total of 132 participants were randomly selected, with half from the top and half from the bottom of the BP-LVMI ranking distribution. These participants were further categorized into five groups (i) low BP+low LVMI: SBP Percentile (PCT) <80^th^ percentile, LVMI:22.35-27.56 gm/m^2.7^; n=10 (ii) high-BP+high-LVMI: SBP PCT≥90^th^ percentile, LVMI: 39.55-44.86 gm/m^2.7^; n=10, (iii) Low BP: SBP PCT <80^th^ percentile, LVMI<36.5 gm/m^2.7^,n=55), and the remaining participants were segregated based on solely on their blood pressure as (iv) mid-BP: SBP PCT ≥80^th^ and <90^th^ percentiles, n=22 and (v) high-BP: SBP PCT ≥90^th^ percentile, n=35.^73^ A complete description of the study approach and their clinical parameters is represented in Supplemental Figure SI, Table 1, and Table 2, respectively, to use high-BP and high-LVMI as the variables. Blood samples and serum were collected to investigate circulatory molecular changes at genetic, epigenetic, and proteomics levels that might influence the development of CV-TOI in hypertensive youth. We excluded youth with symptomatic severe hypertension, on antihypertensive or lipid-lowering medication in the past 6 months, with diabetes mellitus (type 1 or 2), kidney disease, or other known chronic medical conditions. All study participants and their parents provided written informed consent and assent.

### Clinic BP Determination

BP was obtained in the right arm by auscultation with an aneroid sphygmomanometer (Mabis MedicKit 5, Mabis Healthcare, Waukegan, IL) after confirming the correct cuff size for the arm measurement.

After five minutes of rest, four BP measurements were taken at 30-second intervals using the first Korotkoff sound for systolic BP (SBP) and the fifth sound for diastolic BP (DBP). The average of the last three BP measurements was recorded at each visit. Overall, we measured BP with the mentioned auscultatory technique and averaged six BPs taken over two visits to calculate the mean BP. All anthropometric and BP readings were recorded by the same observer. The 2017 Clinical Practice Guideline on BP in youth^74^ was used to determine BP percentile for classification of the participants as low-risk BP (L=mean SBP percentile <80^th^ percentile); mid-risk BP (M=mean SBP percentile ≥80^th^ and <90^th^ percentiles); or high-risk BP (H=mean SBP percentile ≥90^th^ percentile). These thresholds for recruitment were based on our hypothesis that LVH may be found at BP levels lower than 95^th^%. Therefore, we concentrated on recruiting youth with an SBP% above the 80^th^ percentile and above. Anthropometrics and the first set of BP measures were obtained at the first visit and the echocardiogram and second set of BP measures were obtained at the second visit within one month unless a protocol waiver was given for a specific reason. All anthropometric and BP readings were recorded by the same observer.

### Ambulatory BP Measurement (ABPM)

Ambulatory BP is measured with an oscillometric device (SpaceLabs OnTrak, Spacelabs Healthcare, Issaquah, WA) as described in detail previously.^10^

### Echocardiography LVMI Measurement

A standard echocardiogram including two-dimensional-guided M-mode measurement of the LV was obtained for measurement of LV dimensions for calculation of LV mass (LVM). All sites were trained to perform echocardiograms uniformly according to a rigorous protocol from the Cincinnati site. All measurements were performed by a trained sonographer at the Cincinnati site using Cardiology Analysis System (Digisonics, Houston, TX) according to published guidelines.^75,76^ LVM was calculated using the Devereux equation,^77^ and the LVMI was calculated to adjust for differences in body size according to the method of de Simone (LVMI=LVM/ht^2.7^).^78^ The prevalence of LVH was determined using the pediatric cut point for adolescents (38.6 g/m^2.7^)^78^ which relates to the 95^th^ percentile of LVMI.^73^ The mentioned cut-point was selected as TOI in youth as discussed elsewhere.^12^

### Diastolic Function Evaluation

Diastolic function was evaluated measured by evaluating traditional Doppler and Tissue Doppler velocities across the mitral valve from the apical four-chamber view (E/A, E/e’, e’/a’) as described previously.^10^

### Systolic Function Evaluation

Systolic function was measured by tracing the endocardium from the four-chamber view at peak systole and end-diastole using TOMTEC software (TOMTEC Corporation, Chicago, IL) offline to estimate global longitudinal strain (GLS), determine strain rate, and calculate LV ejection fraction (LVEF).^79^

### Arterial Stiffness Evaluation

Pulse wave velocity (PWV) assessment as a measure for vascular/arterial stiffness, another type of hypertensive TOI, is measured using the SphygmoCor CPV System (AtCor Medical, Sydney, Australia) as described previously.^10,80^

### RNA-seq Analyses

Peripheral Blood samples (2.5 ml) were drawn from each volunteer in a PAXgene tube (Qiagen cat# 762165) followed by total RNA isolation using an RNA isolation kit (Qiagen cat# 763134) according to the manufacturer instruction. Total RNA concentration and quality were evaluated using a nanodrop system (ThermoFisher Scientific). RNA samples were subjected to QC and 85 RNA samples (low blood pressure ( Low BP; n=42), elevated/mid blood pressure (mid-BP; n=13), high blood pressure (high-BP; n=30) were sequenced using DNBseq technology at BGI facility, and 20 samples (low blood pressure with low-LVMI (low-BP+low-LVMI; n=10), high blood pressure with high-LVMI (high-BP+high-LVMI; n=10) were sequenced in Illumina HiSeq platforms at Loyola sequencing Core facility, respectively.

Briefly, adapter sequences were removed, and low-quality reads were trimmed using Cutadapt (v. 1.11). The resulting reads were mapped to the human reference genome from Ensembl, GRCh38, using Bowtie2 (v. 2.1.0). The aligned sequencing reads, and a list of genomic features was used as input for the Python package HTSeq (v. 0.6.1p1) to count the mapped genes and generate a table of raw counts. The DESeq2 package (v. 1.14.1) was used to determine differential expression between sample groups using the raw count table by fitting the negative binomial generalized linear model for each gene and then using the Wald test for significance testing. Count outliers were detected using Cook’s distance and were removed from further analysis. The Wald test p-values from the subset of genes that passed an independent filtering step were then adjusted for multiple testing using the Benjamin-Hochburg procedure.

DEseq2 data were compared to identify the common gene signatures upregulated or downregulated among the groups and subjected to functional enrichment analysis using Metascape.^81^ Also their association with hypertension and cardiovascular diseases were identified using DisGeNET database (http://www.disgenet.org/search).^82^ FPKM of commonly dysregulated genes for individual samples were used to correlate its expression with their respective blood pressure and left ventricular mass index using Spearman correlation analysis. Also, FPKM values and normalized counts were used in analyzing the expression levels of anti-angiogenic genes among Mid-BP, high-BP and high-BP+high-LVMI groups.

### miRNA-Seq Analyses

Peripheral Blood samples (2.5 ml) were drawn from each volunteer in a PAXgene tube (Qiagen cat# 762165) followed by total RNA (including miRNA) isolation using a commercial kit (Qiagen cat# 763134) according to the manufacturer’s instruction. Total RNA concentration and quality were evaluated using a nanodrop system (ThermoFisher Scientific). RNA samples were subjected to QC and 109 RNA samples (low blood pressure ( low BP; n=54), elevated/mid blood pressure (mid-BP; n=18), high blood pressure (high-BP; n=37) were sequenced using DNBseq at BGI facility and 20 samples (low Blood pressure with low LVMI (low BP+low LVMI; n=10), and high Blood pressure with high-LVMI (high-BP+high-LVMI; n=10) using Illumina HiSeq platforms at Loyola sequencing Core facility respectively. The data generated from both platforms were processed and analyzed separately using standard bioinformatic pipelines outlined in the above method section, and differential expression was analyzed using DESeq2.

Briefly, the adapter sequences were removed, and low-quality reads were trimmed using Cutadapt (v. 1.11). The resulting reads were mapped to the human reference genome from Ensembl, GRCh38, using Bowtie2 (v. 2.1.0). The aligned sequencing reads, and a list of mature miRNA genomic coordinates from miRBase (v. 21) were used as input for the Python package HTSeq (v. 0.6.1p1) to count the mapped miRNAs and generate a table of raw counts. The DESeq2 package (v. 1.14.1) was used to determine differential expression between sample groups using the raw count table by fitting the negative binomial generalized linear model for each miRNA and then using the Wald test for significance testing. Count outliers were detected using Cook’s distance and were removed from further analysis. The Wald test p-values from the subset of genes that passed an independent filtering step were then adjusted for multiple testing using the Benjamin-Hochburg procedure. DEseq2 data were compared to identify the common miRNA signatures upregulated or downregulated among the groups. TPM of selected miRNAs predicted to target *VASH1* 3’UTR for individual samples were used to correlate its expression with their respective blood pressure and left ventricular mass index using Spearman correlation analysis.

### Whole-genome Bisulphite DNA Seq Analyses

Blood samples (2.5 ml) were drawn from each volunteer in a PAXgene tube (Qiagen cat# 762165), followed by genomic DNA isolation using a DNA isolation kit (Qiagen cat# 80204) according to the manufacturer’s instructions. DNA sample concentration and quality were evaluated using a nanodrop system (ThermoFisher Scientific). DNA samples extracted from Low BP (n=10), Mid-BP (n=10) and high-BP+high-LVMI (n=10) groups were processed for global DNA methylation analysis via the whole-genome DNA methylation approach in DNBseq-G400 at BGI facility. A series of pairwise comparisons of methylated sites were made to identify differential methylation regions. Specifically, we compared the methylated sites determined in the Mid-BP group with the Low BP group, resulting in 10 pair-wise comparisons. Similarly, we compared the methylated sites determined in the high-BP+high-LVMI group with the Low BP group, resulting in another 10 pairwise comparisons. Putative DMRs were identified by comparison of the sample1 and sample2 methylomes using windows that contained at least 5 CpG sites with a 2-fold change in Methylation level and Fisher test p-value <=0.05. The following formula was used: degree of difference=log2 Rm1/log2Rm2; Rm1 and Rm2 represent the Methylation level of methyl-cytosine for sample1 and sample2, respectively. 0.001 will replace Rm1 (or Rm2) while it is 0.^83^ The DMR regions and associated genes thus identified were subjected to functional enrichment analysis using Metascape.^81^ Next, the DEG list from the RNA sequencing data was compared with that of the identified DMR gene list to correlate the methylation pattern and its expression level.

### Mass Spectrometry

Serum samples (1 ml) were drawn from each volunteer in assigned tubes (BD bioscience cat#), followed by the addition of a protease inhibitor cocktail (ThermoFisher Scientific cat# 78430) according to the manufacturer’s instructions. Also, the concentration and quality of protein samples were evaluated using Bradford assay, and samples were processed for proteomic analysis via mass spectrometry at Cornell University proteomics core, USA. The 30 serum samples (low BP =10, mid-BP=10, high-BP high-LVMI=10) were randomly grouped into three sets consisting of samples from all three groups. Every set has a pool of all 30 samples, and its intensity is used to normalize each sample’s abundance intensity. Results were received as a protein abundance for ∼298 protein list that was further analyzed at our lab to identify the differentially expressed proteins between the groups based on their relative ratio. The identified DEPs were subjected to functional enrichment analysis using Metascape.^81^

### Statistical Analyses

Analyses of clinical data were conducted using SAS (version 9.4, SAS Institute Inc., Cary, NC). *DESeq2* method employing WALD test was used in identifying significant differential expression of mRNA and miRNA. We use P ≤ 0.05 and the effect size of Log2Fold Change ≥ 0.5 as the default threshold to judge the magnitude of the gene expression difference between groups. Fisher t-test was used in identifying significant differentially methylated regions and students t-test, P <0.05 for the abundance of protein levels among the groups. Wherever needed, we used the following to account for multiplicity-multiple unpaired t-test was performed for group analysis, in which multiple comparisons using the Holm-Sidak method with P value (adjusted P Value) threshold of 0.05 to compare the differences among the groups. Analyses were conducted using Prism 10.1.2 (GraphPad Software, LLC) wherever needed, such as the Spearman correlation of SBP PCT with the FPKM values of individual genes or TPM values of miRNAs. To compute correlations, Spearman correlation coefficients were used to specify the significant correlation using p-value <0.05 calculated using two-tailed analysis. The graphical approach was undertaken for further analysis and presentation of data by using the GraphPad Prism 10.1.2 and FunRich program. Additional details of the analyses used are included in the figure captions.

## Data Availability

The high throughput sequencing data (raw and processed) are deposited in the Gene Expression Omnibus (GEO) database and available with accession number GSE262821 as a super series. The accession number for sub-series are GSE262828 (bulk RNA Seq 1), GSE262814 (bulk RNA Seq 2), GSE262815 (bulk RNA Seq 3), GSE262818 (small RNA Seq 1), GSE262819 (small RNA Seq 2) and GSE262817 (Methylation Bisulfite Seq). The mass spectrometry proteomics data have been deposited to the ProteomeXchange Consortium via the PRIDE partner repository with the dataset identifier PXD050915.

## Acknowledgments

Dr. Urbina, Dr. Becker, and Dr. Sadayappan were supported by AHA Strategic Focused Network Funding (SFRN23680000). Dr. Urbina was supported by NIH UL1 TR001425 (CTSA). Dr. Sadayappan has received support from National Institutes of Health grants (R01 AR079435, R01 AR079477, R01 HL130356, R01 HL105826, R01 AR078001, and R01 HL143490), the American Heart Association, Institutional Undergraduate Student (19UFEL34380251), Transformation (19TPA34830084 and 945748) awards and the PLN Foundation (PLN crazy idea) awards. We thank the Proteomic and Metabolomics Facility of Cornell University for generating the mass spectrometry data.

## Non-standard Abbreviations and Acronyms

BP: Blood pressure
CV: Cardiovascular
DEG: Differentially expressed gene
DEP: Differentially expressed protein
DMR: Differentially methylated region
ECM: Extracellular matrix
FDR: False Discovery Rate
FPKM: Fragments per kilobase of exon per million
GO: Gene Ontology
IGF: Insulin growth factor
KEGG: Kyoto encyclopedia of genes and genomes
LVEF: Left ventricular ejection fraction
LVFS: Left ventricular fraction shortening
LVH: Left ventricular hypertrophy
LVID;d: Left ventricular internal diameter end-diastole
LVID;s: Left ventricular internal diameter end-systole
LVMI: Left ventricular mass index
miR: microRNA
PBMCs: Peripheral blood mononuclear cells
PH: Primary hypertension
SBP PCT: Systolic blood pressure percentile
TPM: Transcripts per million
TOI: Target organ injury
VASH1: Vasohibin-1
VEGF: Vascular endothelial growth factor

---

**Supplemental Figure S1.**
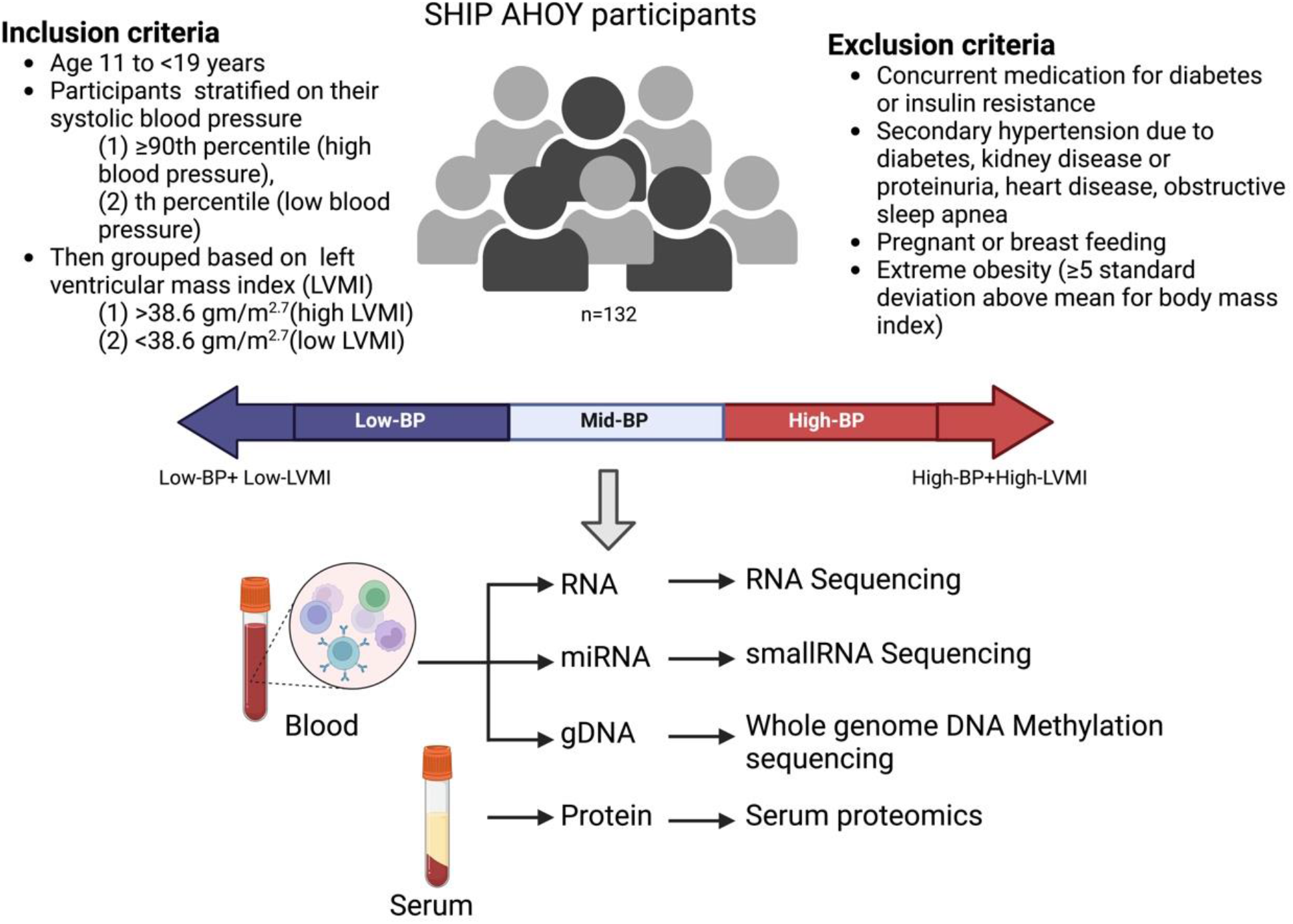
Human study experimental approach. This schematic depicts the methodology and study approach in the present study. We examined a SHIP AHOY cohort of youths with the indicated inclusion and exclusion criteria. The study participants were stratified based on their blood pressure and 132 adolescents (mean age ∼15 years, male-57.58%) were included for this cross-sectional study (Table 1, Methods section) and grouped based on their left ventricular mass index (Table 2, Methods section). Blood and serum samples collected from the participants were utilized to isolate total RNA, miRNA, genomic DNA and protein samples that were later utilized to perform RNA sequencing, miRNA sequencing, whole-genome DNA methylation sequencing and serum proteomics analyses. The sequencing and proteomics data were further analyzed and correlated to identify the candidate genes and pathways involved in the early cardiovascular target organ injury (Figure Created with BioRender.com). **gDNA**: genomic DNA, **miRNA:** microRNA.

**Supplemental Figure S2.**
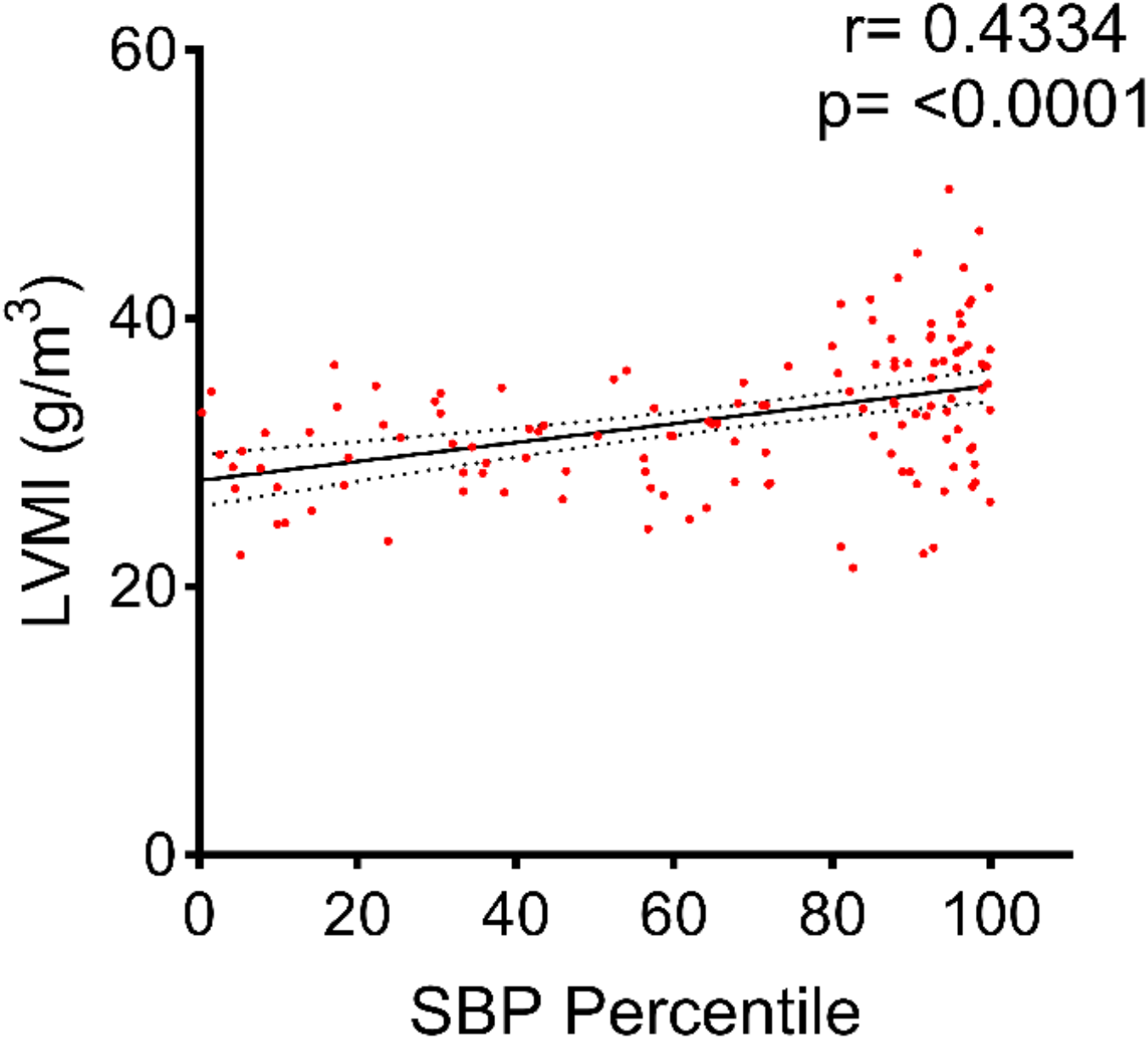
Correlation of LVMI with systolic blood pressure percentile in adolescents of SHIP AHOY participants. Each data point represents an individual participant’s SBP Percentile values against their LVMI (n=132). The Spearman correlation coefficient and its significant value are 0.4334 and p<0.0001 respectively, indicating a significant positive correlation between SBP percentile and LVMI. LVMI: Left Ventricular Mass Index; SBP: systolic blood pressure.

